# Breathing in sync: how a social behavior structures respiratory epidemic risk in bottlenose dolphins

**DOI:** 10.1101/2023.12.01.569646

**Authors:** Melissa A. Collier, Ann-Marie Jacoby, Vivienne Foroughirad, Eric M. Patterson, Ewa Krzyszczyk, Megan Wallen, Madison Miketa, Caitlin Karniski, Sarah Wilkin, Janet Mann, Shweta Bansal

## Abstract

Dolphin morbillivirus has caused mass mortalities in dolphin populations globally. Given their role as ecosystem sentinels, mass mortalities among these populations can be detrimental. Morbillivirus is transmitted through respiratory droplets and occurs when dolphins breathe synchronously, a variable social behavior. To assess the role of variable social behavior on disease risk empirically, we collected behavioral data from two wild bottlenose dolphins populations (*Tursiops* spp.), developed network models that synthesize transmission contacts, and used an epidemiological model to predict disease consequences. We find that juveniles have more contacts than adults, adult males have more contacts than adult females, and that individuals preferentially contact others in their own demographic group. These patterns translate to higher infection risk for juveniles and adult males, which we support using data from a morbillivirus outbreak. Our work characterizes the impact of bottlenose dolphin social dynamics on infectious disease risk and informs the structure of vulnerability for future epizootics.

## 1.0 Introduction

The world’s oceans are changing rapidly from anthropogenic disturbances and human-induced climate change^1,2^. Already, these changes have included major effects on the emergence and spread of infectious diseases. For example, warming ocean temperatures have increased the growth rates and habitats of pathogens and broadened their repertoire of potential hosts^3,4^, and high levels of shipping traffic and shore runoff have facilitated the emergence of pathogens in the ocean^5,6^. Facing these threats on the front lines are delphinid species (i.e., dolphins), ocean predators and sentinel species that are essential for healthy ocean ecosystems^7–9^ but among whom disease and mass mortality have surged in recent decades^1,10^. Specifically, catastrophic die-offs of dolphins caused by the infectious pathogen dolphin morbillivirus (DMV) are a global phenomenon^11^. Our work aims to understand the underlying drivers of disease spread in dolphin populations so that we might be able to more effectively and efficiently estimate infection outcomes for these essential species^12^.

The most recent mass mortality due to DMV occurred in July of 2013 in bottlenose dolphins (*Tursiops* spp.) along the US Atlantic coast in which more than 1,650 dolphins were recovered dead over a two-year period. The bottlenose dolphins in this area of the northwest Atlantic likely comprise animals from multiple populations^13^. Some of these populations are resident to restricted areas, but other “coastal” populations exhibit large seasonal migratory movement. The two coastal bottlenose dolphin populations along the northwestern Atlantic are considered ‘depleted’ (or under optimal population levels) by the National Marine Fisheries Service because of disease mortality^13,14^, indicating that mass die-offs among dolphin species are detrimental to their populations and ecosystems. Indeed, it is estimated that the abundance of these coastal populations declined by more than 40% during the 2013 DMV outbreak, which is now the deadliest known DMV epizootic^15,16^.

Life history factors are often linked to mortality in DMV epizootics. For example, in a 2007 Mediterranean epizootic a greater proportion of juveniles were found dead compared to adults^17^, suggesting that infection outcomes differ across age classes. Numerous factors may drive differentiation in infection outcomes across individuals, such as host immunity^18^, or environmental and anthropogenic conditions^19^. What is more poorly understood is the role of social behavior across demographic groups in structuring infection risk^20,21^. Highly social cetaceans such as bottlenose dolphins live in fission-fusion societies where members maintain long-term preferential bonds, but exhibit associations that change in time and space^22,23^. Stable affiliations include mother-calf bonds^24^, and long-term alliances formed among males^25^, while various social phenotypes can be found among juveniles and females^24,26,27^. This stability of strong long-term bonds coupled with social heterogeneity leads us to hypothesize that social structure could drive disease variability and severity in bottlenose dolphins and delphinids with similar social structures.

In order to study disease transmission and predict epidemic consequences based on social behavior, it is important to identify the specific behaviors that allow for infection propagation between an infected and susceptible individual^28^. For example, since DMV is a respiratory transmitted virus, transmission in cetaceans likely occurs by synchronized breathing, defined as two or more individuals surfacing and breathing simultaneously (e.g., within two seconds) and in close proximity (e.g., within two meters). The expired “blow” that is forcefully released from their blowholes is a combination of mucus, water and air, and is thought to be sufficient for droplet transmission of respiratory viruses^11^. Synchronized breathing occurs in a variety of contexts in delphinid species, is important for both socialization and parental care^29–32^, and is expected to be heterogeneous across both demographic (e.g., age, sex)^33^, and behavioral states (e.g., foraging, consorting)^25,34,35^.

Contact networks are often used by researchers to capture heterogeneity in transmission behaviors and to provide a tractable method to represent variation from social and demographic structure^20,36^. With the contact structure of the population represented in a network model, an epidemiological modeling approach can be an ethical and economical way to test hypotheses regarding potential factors that drive disease spread in wildlife and can play a key role in predicting the risk of epidemics in wild marine populations^37,38^. Few such studies exist for DMV. In fact the majority of marine mammal disease studies focus on the spread of the similar phocine distemper virus (PDV) in seals^39–44^. There has been only one study^15^ that examines the transmission dynamics of DMV in the western North Atlantic bottlenose dolphin populations, which focuses primarily on large-scale spatial dynamics of the epizootic and does not explicitly consider contact dynamics between individuals. However, this study was able to estimate important features of DMV including the infectious period (5-10 days) and the basic reproduction number, R0 (0.9-2.6)^15^.

Motivated by the devastating consequences of the 2013 western North Atlantic DMV epizootic and the urgent need for data-driven work in delphinid populations, we leverage a study of wild bottlenose dolphins (recently classified as *Tursiops erebennus*^45^) that 1) utilize the region where the plurality of 2013 DMV deaths occurred (fig. S22), and 2) are currently below optimal sustainable population levels from disease outbreaks^14^. We hypothesize that infection risk will vary based on individual differences in synchronized breathing contact among age and sex classes which could impact population level vulnerability. With this work, we aim to characterize the impact of dolphin social structure and dynamics on disease risk using empirical data, and inform future data collection on behavior and disease. A better understanding of disease transmission in this complex marine species may also allow us to develop strategies that minimize the detrimental effects of perturbations to this vulnerable population and to the overall ecosystem.

## 2.0 Results

Our central hypothesis is that social structure drives disease dynamics in bottlenose dolphins (*Tursiops* spp.). To assess this empirically, we collected data from a wild dolphin research site in the Potomac-Chesapeake Bay region, USA (PC). Since it is not feasible to observe complete contact networks, we collected data and estimated 1) the average number of synchrony contacts over a DMV infectious period, *μ*, for the following demographic groups, *g*: adult males, adult females and calves, and juveniles (synchrony degree, *k*_*g*_(*μ*)). We then estimated the degree of mixing among these groups. We developed a generative network model to construct realistic contact networks informed by these empirical data. To predict disease consequences, we used an epidemiological model on these generated contact networks and examined the epidemic consequences across age and sex classes ^20^. We test the generalizability of our behavioral findings by comparing to empirical data from a second long-term study of wild bottlenose dolphins (*Tursiops aduncus*) from Shark Bay, Australia (SB). Finally, to support our disease models, we compared our results to National Marine Fisheries Service mortality data from the 2013 DMV epidemic in bottlenose dolphins (*Tursiops* spp.) along the western North Atlantic. For all results, uncertainty is represented by standard deviation. The specifics of our methods are summarized in Fig. 1, with a more detailed summary in fig. S1.

**Fig. 1:**
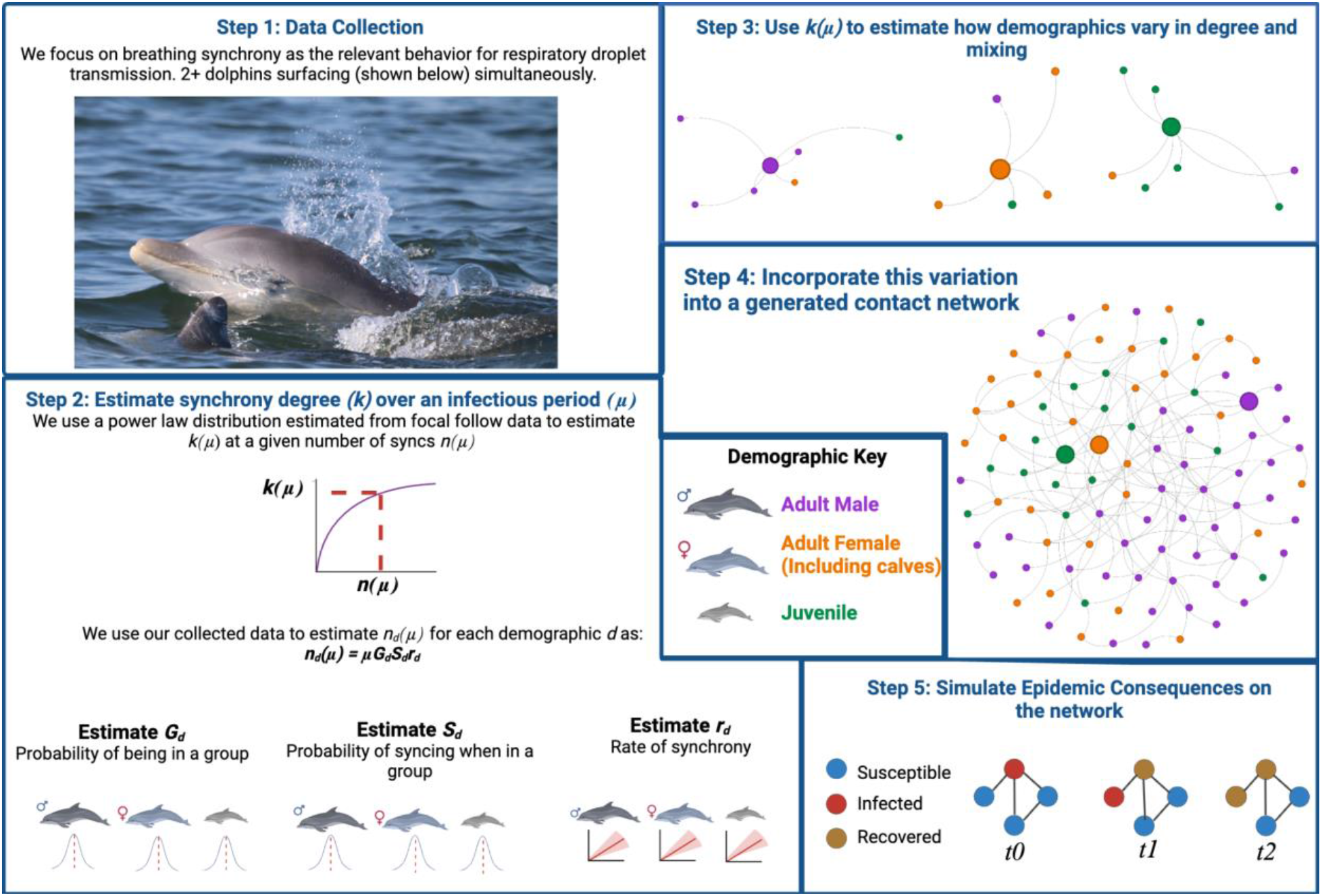
Summary of methods. We first collected data on synchronized breathing behavior on bottlenose dolphins to estimate how respiratory droplets might transmit disease in cetaceans (Step 1). We then use the data to estimate the synchronized breathing behavior over the course of a DMV infectious period (Step 2) and use these results to model a contact network that is representative of synchronized breathing and its variation across individuals (Steps 3-4). Finally we simulate the spread of disease across this contact network to characterize epidemic outcomes (Step 5). Photo credit: Potomac Chesapeake Dolphin Project, NMFS Permit No. 19403. Created with BioRender.com

### 2.1 Bottlenose dolphin synchronized breathing is structured demographically

We generated estimated degree distributions and demographic mixing matrices for the PC bottlenose dolphins to represent synchronized breathing contact over a DMV infectious period. We found that synchrony degree, *k*_*d*_(*μ*), and the variation in degree varies significantly across demographic groups for a DMV infectious period (F = 1152; d.f. = 2; p <0.001) (Fig. 2A (right) and 2B). Juveniles had the highest average degree (*k*_*juv*_(*μ*)= 36.3), followed by adult males (*k*_*male*_(*μ*)= 30.8), while adult females and calves had the lowest (*k*_*fem*_ (*μ*)= 21.9). We found that our estimate of demographic-specific degree structure over an infectious period is also supported by our empirical measurement of demographic-specific degree (Fig. 2A, left) where juveniles have a significantly higher observed degree than adult females (p= 0.008), though the higher degree of males compared to adult females is not statistically significant (p= 0.295).

**Fig. 2:**
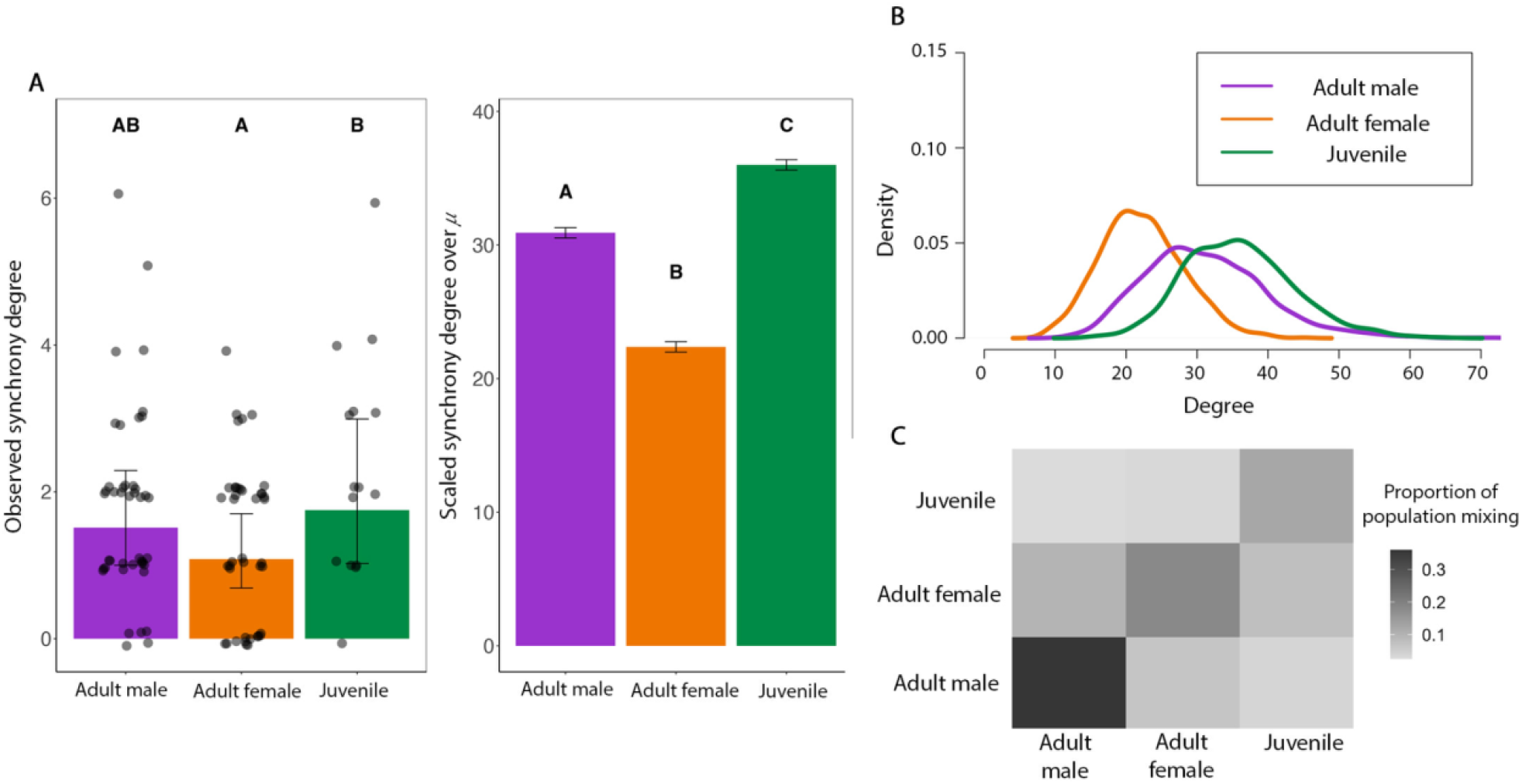
The degree distributions and mixing matrices generated from PC data. A (left), shows the average observed sync degree over the course of an average PC focal follow (n=99) using a GLMM to control for different variables where points show the raw data, letters indicate significant differences, error bars represent standard error from the mean. A (right) shows how this observed degree scales up to an average DMV infectious period for n=1000 estimates for each demographic group using our model from Fig. 1. Letters represent significant differences based on a one-way ANOVA test and error bars represent standard deviation. We used these data to estimate (B) a demographic specific degree distribution and (C) a demographic mixing matrix, where each cell represented the proportion of all observed synchrony dyads that existed between a focal demographic (row) and each demographic group (columns) (termed proportion of population mixing) for the PC over a DMV infectious period.

These dolphins are also significantly more assortative in their synchronized breathing interactions than what would be expected at random (Newman’s assortativity=0.47; z score = −5.09; p = <0.0001), suggesting that individuals breathe synchronously most often with individuals of their own demographic (Fig. 2C). Indeed, we saw that our predicted mixing values and confidence intervals were higher along the diagonal of our mixing matrices (where assortative mixing is represented) compared to the expected values for all three demographic groups (fig. S11).

For testing generalizability, we compared these results to contact structure in the SB population and found consistent patterns (F = 1788; d.f. = 2; p <0.001) : adult females and mother calf pairs had the lowest synchrony degree (*k*_*fem*_(*μ*)= 6.4) followed by juveniles (*k*_*juv*_ (*μ*)= 10.5) and adult males (*k*_*male*_(*μ*)= 12.2) (fig. S9, S10); and synchrony degree was more assortative than expected at random (Newman’s assortativity =0.39; z score = −8.65; p = <0.0001) (fig. S12).

### 2.2 PC dolphins have higher infection risks than SB dolphins

We considered the spread of disease on networks representative of demographic degree and assortativity for bottlenose dolphins. The average reproductive number, *R*_*0*_ experienced by a host population during an outbreak is a function of the population’s contact structure and the transmissibility of the pathogen. We found that that in the PC dolphins *R*_*0*_ = *1*.*9*, which is the average reproductive number observed for past morbillivirus outbreaks in marine mammal populations, was achieved by a weakly transmissible pathogen with transmissibility, *T* = *0*.*055*. To achieve the same *R*_*0*_ in the SB population required a moderately transmissible pathogen of *T* = *0*.*135* (fig. S16). With these transmissibility values, we simulated epidemics representative of DMV spread in dolphin populations. We found that this resulted in about 70% of the total population infected and with an approximately 70% likelihood of an epidemic occurring after a random introduction of disease into the population (fig. S17). Additionally, our sensitivity analysis showed our results are robust to potential errors in demographic group assignment (fig. S18). For context, population abundance estimates place the 2013-2015 DMV outbreak in certain Atlantic bottlenose dolphin populations as resulting in a 40% population decline^46^, suggesting a high case fatality rate or substantial indirect loss of life due to the outbreak.

### 2.3 Infection risk is age- and sex-structured

We simulated the spread of disease on networks representative of demographic degree and assortativity. We calculated the proportion of each demographic group that was infected across all networks and simulations for which an epidemic occurred. We found that our models predicted juveniles to be disproportionately infected in DMV epidemics compared to adults (t = 82.353; df = 42.068; p-value < 0.0001) (Fig. 3A). Within adults, males were predicted to be significantly more at risk for infection than females (t = 91.13; df = 46.903; p-value < 0.0001) (Fig. 3D). To compare to epidemic outcomes in the SB population, we performed the same epidemic simulation experiments, and found highly consistent patterns of transmission (juveniles more at risk for infection than adults: t = 14.283; df = 32.522; p-value < 0.0001; adult males more at risk for infection than adult females: t = 161.2; df = 46.6; p-value < 0.0001) (Fig. 3B and 3E).

**Fig. 3:**
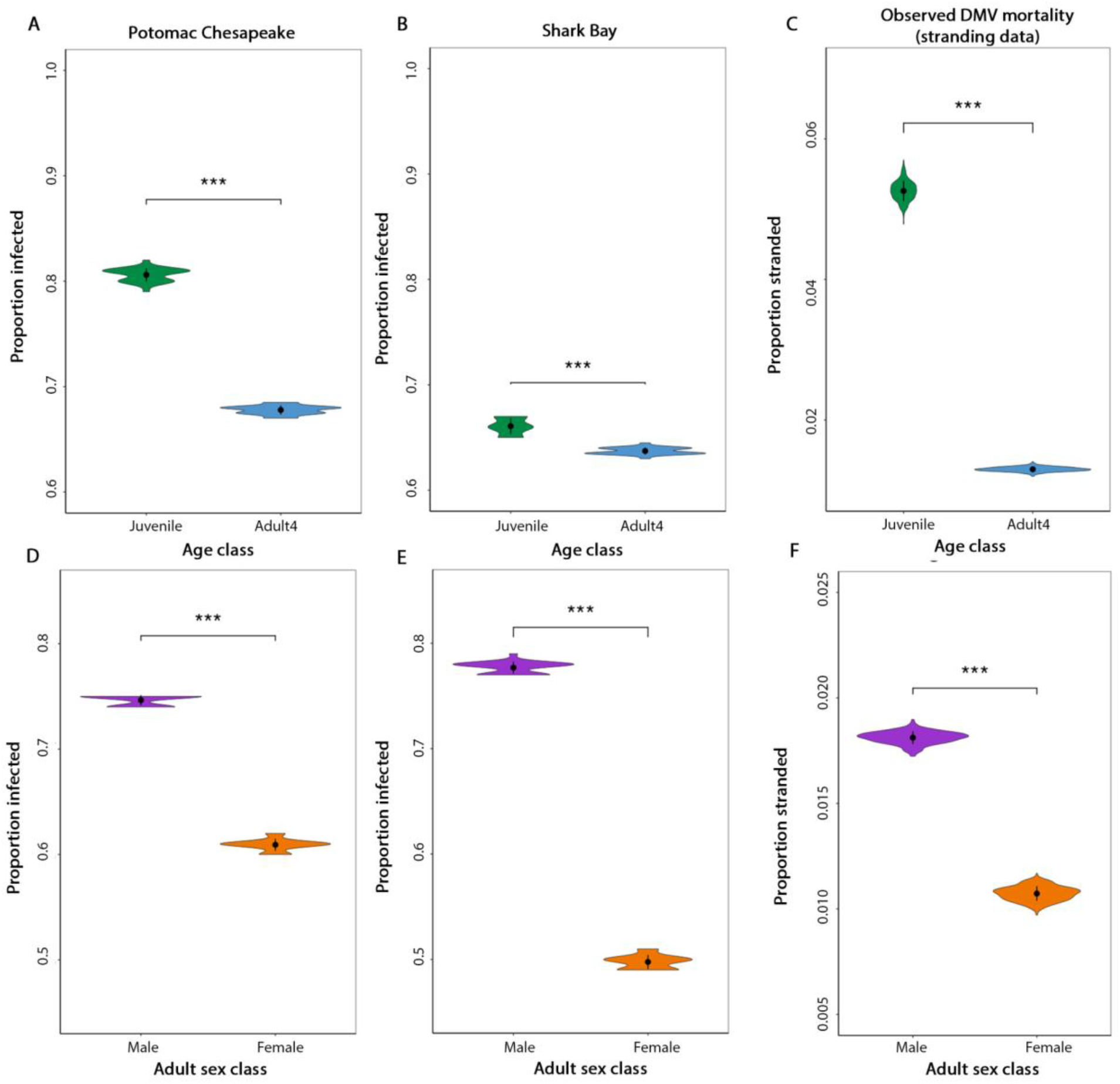
Epidemiological model results compared to the stranding data. The top row shows the model results for the proportion of each age class infected in the PC (A) and SB (B) based on 100 disease simulations on 25 network models for each population. (C) shows the estimated proportion of each age class that was recovered during the 2013 DMV epizootic on the Atlantic Coast, derived from the stranding data (n=504 strandings). The bottom row shows the proportion of each adult sex class that was infected in our models in the PC (D) and SB (E) based on 100 disease simulations on 25 network models for each population. (F) shows the proportion of adults of each sex class recovered on the Atlantic Coast, derived from the mortality data (n = 242 strandings). Differences were significant across all comparisons (asterisks) based on a t-test, and uncertainty represents standard deviation.

We show support for our results with empirical mortality data from strandings that occurred during the Atlantic DMV outbreak of 2013-2015. We found that the demographic biases in infection predicted by our model results were reflected in the observed stranding data with juveniles stranded at significantly higher rates than adults when controlling for the estimated population size for each age class (t = 554.43; df = 540.51; p-value < 0.0001) (Fig. 3C and Fig. S14), and adult males stranded at higher rates than adult females (t = 343.04,;df = 995.92; p-value < 0.0001) (Fig. 3F and fig. S14).

Thus, our results showed age- and sex-specific biases in infection risk. To better understand whether this risk bias is structured by demographic-specific degree distributions (the average and variation in the number of contacts per demographic group) or demographic assortativity (the tendency of demographic groups to mix more within groups), we ran epidemic experiments on null models. We found that in a population with only demographic-specific degree structure, epidemic sizes (the proportion of the total population infected) were significantly lower (PC: F=8504, d.f. = 2, p<0.0001; SB: F=1025, d.f. =2, p <0.0001) (fig. S19). Importantly, we also found that with only demographic assortativity present, the demographic biases in infection outcomes no longer existed (PC: F=2.672, d.f.=2, p= 0.076; SB: F =0.437, d.f =2, p = 0.648) (fig. S20). This suggests that degree is the most important driver of the demographic-specific infection risks and assortative mixing drives the overall population infection burden.

### 2.4 Population infection risks are amplified when disease is introduced by adult males or juveniles

We examined how population infection burden differs depending on which demographic group begins the infection in our 25 PC network models. We found that when infection is introduced to the population by a juvenile, disease burden was higher for both age classes compared to when infection is introduced by an adult (F = 67.15, d.f =3, p <0.0001) (Fig. 4, left). Similarly, when the index case is an adult male, we saw that both adult males and adult females had higher infection burdens than when the index case is an adult female (F =168.3, d.f. =3, p <0.0001) (Fig. 4, right). This suggests that demographically structured social behavior not only drives infection risk within demographic groups, but also epidemic risk at the population level.

**Fig. 4:**
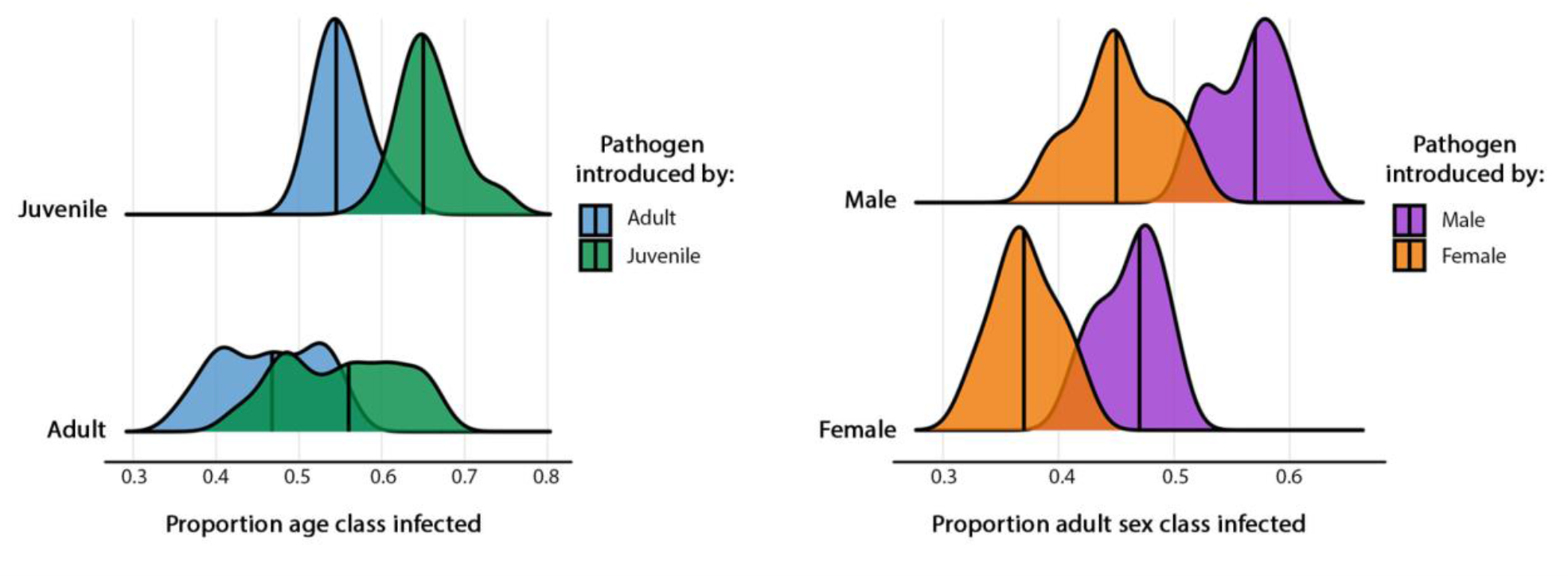
The effect of pathogen introduction on epidemic outcomes. The proportion of each age class (left) and adult sex class (right) that is infected in our epidemiological model for the PC, based on in what demographic group the infection began or was introduced (colors) for n=100 disease simulations on 25 PC network models. Uncertainty represents standard deviation.

## 3.0 Discussion

Our analysis shows that there are significant differences in synchronized breathing contact dynamics across age and sex in bottlenose dolphins. While infection risk can be structured demographically due to a variety of other factors such as environmental conditions or host immunity^47,48^, our goal in this work is to present a parsimonious explanation for observed heterogeneity in infection risk based on social behavior only.

Our comparative analysis of two wild bottlenose dolphin studies, one in the heavily impacted waters of an Atlantic Ocean bay^49^ and the other in the protected waters of an Indian Ocean bay, highlights that relative infection risk is structured similarly despite notable ecological differences between the two research sites. However, we also demonstrate a substantially higher absolute risk of epidemics occurring in PC dolphins relative to the SB population, an idea supported by the fact that certain populations of the Atlantic bottlenose dolphins have decreased by 40-50% twice in 25 years from morbillivirus^14,46,50^, while SB has no recorded mass mortality events from infectious diseases. Our work explains differences in vulnerability based on the fundamental social behavior of synchronized breathing, and quantifies the risk in terms of a higher number of and more homophilic transmission contacts in the PC. The substantial difference in social connectivity can be explained by a number of differences. First, the dolphins in the PC are thought to be a part of at least three different populations of Atlantic bottlenose dolphins^13^, meaning there are likely more individuals (∼10,000) compared to SB (∼2,700)^51^. Indeed, the average group size for SB based on our data was 3.7 compared to 18.7 in the PC. Second, the SB population is a year-round, residential population while the animals in the PC are considered seasonal and migratory^13,52–54^. These factors drive notable changes in the observed activity budgets of individuals, with the SB population spending ∼35% of their time foraging (a largely solitary activity^23,55^) and ∼22% time traveling ^56^, while the PC preliminary estimates suggest individuals spend a large majority of their time traveling (a largely social activity) when present at the PC study area (∼7 months). While it is possible that activity budgets for PC dolphins might change in other parts of their habitat that we have not observed, we suggest that for at least half the year their contact rates might be elevated due to long times spent traveling. Our comparative approach thus highlights consistencies in demographic contact patterns and disease risk structure despite ecological differences, and generates hypotheses about the role of metapopulation structure and seasonal changes in distribution like migration on population vulnerability to threats.

We also find that there is significant assortativity for synchronized breathing among demographic groups. This is consistent with past findings^33^, and suggests a preference for social associations within similar age and sex classes. This effect was stronger in the PC than in SB, which may also contribute to increased infection risk in the former. Increased assortativity by trait (e.g., age or sex) can increase infection burden in networks if the number of contacts are also structured by that trait^57^. Indeed, we found that demographic assortative mixing had a significant impact on epidemic risk in the PC, but less so in SB. It is possible these differences are due to an observation bias in that PC focal follows were conducted over a shorter period of time and a different life-history stage compared to SB focal follows. In adults, male-male synchronized breathing is likely higher during SB breeding seasons when males are consorting females^25^. Synchronized breathing has also been shown to increase with tourist boat traffic^34,58^, as a potential behavioral response to a threat. If both these trends are also the case in the PC, we might see higher rates of assortative mixing due to only observing synchrony interactions during a period of heightened interactions. Future work on the disruptions to contact structure due to seasonal dynamics or anthropogenic factors could reveal important findings for understanding population level infection risk over an annual period.

Our results also highlight that adult females and mother-calf pairs have a lower infection risk than adult males due to their smaller synchrony degree. Synchronized breathing is important for social bonding, particularly in bottlenose dolphin populations where male alliances are common, such as in Shark Bay^25^. In contrast, female associations are considered weaker on average, and vary distinctively across individuals between being almost solitary to having moderate strength relationships with other females^24,26,59,60^. We still know very little about the sex specific behaviors of the dolphins that visit the Potomac-Chesapeake. However, our results show that the differences in synchrony degree among males and females are similar compared to Shark Bay, suggesting that these sex specific behaviors may be maintained across populations and different species of the genus *Tursiops*. Male-biased infection rates due to behavioral differences between males and females is a phenomena seen in many non-human animal species^12,61,62^. Our past work has also demonstrated that social behavior puts male bottlenose dolphins at high infection risk for pathogens of a variety of transmission modes including sexual and skin to skin transmission^21^.

We observed a difference in infection risk between adults and juveniles, suggesting that synchrony degree may be highest in the juvenile period. The relationships formed in the juvenile period are likely critical for survival, forming future social bonds, and reproductive success for both males and females^63–65^. Previous work on juvenile associations in Shark Bay has shown that juveniles socialize significantly more than adult females in the same population, and compared to adults from other bottlenose dolphin populations^27,66^. The higher synchrony degree seen in juveniles compared to adults in the PC reflect these findings; synchronized breathing may be most important as bonds begin to form and less important in the maintenance of stable bonds.

The adult/juvenile effect in SB is not as large, which is likely driven by the high adult male synchrony degree observed in this population which has one of the most complex male alliances in the non-human animal world. Male alliance structure and social bonds have not been investigated in the PC dolphins, but our results suggest that while synchronized breathing is still important for social bonding between adult males, it might be even more important in the juvenile period for these individuals.

We found that disease consequences were worse overall when infections originated in juveniles compared to adults, and in adult males compared to adult females. This suggests that while certain demographic groups are at greater risk of infection because of their high synchrony degree, the entire population is also more vulnerable to infection when infections are initiated in these groups. These results demonstrate the importance of considering how different demographic groups might be more susceptible to contracting novel pathogens, as there could be varying impacts at the population level. The emergence of DMV in the Atlantic bottlenose dolphin populations is thought to be from interactions with pilot whales who are asymptomatic carriers of morbillivirus^11^. Other highly pathogenic viruses including avian influenza have also been recently discovered in marine mammals, including bottlenose dolphins, likely from interactions with sea birds^67,68^. Interactions with humans might also affect the emergence and spread of infectious diseases in dolphins. A recent study found evidence for human pathogens present in bottlenose dolphins that have high human contact due to a provisioning program in Shark Bay, Australia^69^, suggesting opportunity for a direct pathogen transmission from humans to animals. Understanding variation in interspecies (both human and nonhuman) interactions across demographic groups could provide a better understanding of overall population infection risk for current and emerging infectious diseases.

Disease models have been used in the protection of vulnerable, threatened, populations of ecosystem sentinels worldwide, and our work contributes to such research. Our models use empirical data to predict demographic groups that are most at risk during epidemic outbreaks. These predictions can be used to understand downstream impacts on population dynamics and recovery, as well as for developing potential targeted control strategies for reducing disease burden^40^. The control of disease in dolphins remains a challenge that requires effective and long-term solutions. Candidate vaccines against morbillivirus exist for DMV^70^, but the vaccination of free living, obligate marine wildlife, such as dolphins, is fraught with logistical and ethical challenges. Therefore, we recommend focusing on reducing factors that are known to increase disease burden in marine species^19^. Our work can also be expanded to understand how behavior might affect infection risk among overlapping populations of Atlantic dolphins. We suspect that multiple populations of dolphins are seen in the PC study area, but our analysis does not take into account potential differences among them. Future work should therefore examine how this potential heterogeneity in mixing within and among populations might affect the spread of diseases along the US Atlantic Coastline.

## 4.0 Materials and Methods

### 4.1 Data collection

To evaluate contact dynamics in bottlenose dolphins, we used behavioral data collected from boat-based observational surveys and focal animal sampling^56^. When dolphins were sighted, we first conducted a survey consisting of a 5-min scan sample to determine group composition (individual identities determined via photo identification (ID) of unique dorsal fins ^71^), life history information (Supplemental Appendix 1), and predominant group behavioral activity (see ^31^ for behavioral activity definitions). To evaluate synchronized breathing, we then conducted focal follows on an individual dolphin or a mother and her dependent calf (considered a single focal animal due to largely overlapping behavior) observed in the initial survey for a predetermined amount of time between 15 minutes and 3 hours. Focal follows are one-minute interval point samples where we record the behavioral activity of the focal animal. The number of behavioral events (e.g., synchronized breathing) were collected continuously throughout the follow for the focal individual, as well as the ID of their contacts. We used survey and focal follow data from two studies on wild bottlenose dolphins: The Potomac-Chesapeake Dolphin Project (*Tursiops erebennus*) and The Shark Bay Dolphin Research Project (*Tursiops aduncus*).

Our primary populations of interest are those studied by the Potomac-Chesapeake Dolphin Project (PC), a long-term study initiated in 2015 containing detailed data on individual life histories and behavior of bottlenose dolphins that inhabit the waters where the plurality of the mortalities occurred during the 2013 DMV epidemic (fig. S22)^16^. These dolphins are thought to be a part of at least three different populations of Atlantic bottlenose dolphins. For the purpose of this analysis we assume that all the Atlantic bottlenose dolphin populations in the study area have similar contact trends across demography. We collected data over an eight year time period between April and October 2015-2022 in Maryland and Virginia waters from the lower tidal Potomac River (southeast of the Governor Harry W. Nice Memorial Bridge, N 38.361595, W 76.997144) to the middle Chesapeake Bay (specifically Ingram Bay N 37.790233, W 76.997144). We collected 99 focal follows on 97 PC dolphins between June and September 2015-2022 with an average follow time of 25 minutes.

For testing generalizability across bottlenose dolphin populations, we also used data from the Shark Bay Dolphin Research Project (SB), a population that has not yet been documented to have experienced a DMV epidemic. This long-term study was initiated in 1982 and contains more than 35 years of detailed data on individual life histories including sex and age^72^. We used 610 focal follows on 132 SB dolphins conducted over an eight year time period between April and December 2010-2017, with an average follow time of 119 minutes.

For the PC, we used a set of rules for assigning age and sex classes to each focal dolphin and their contacts using sighting records, behavioral data, and photographic data, with high or low confidence (more information in Supplemental Appendix 1). From these rules, we determined the following demographic groups: juvenile (n=15), adult female and mother-calf pairs (n=41), and adult male (n=43). Given our sex assignment rules and juvenile behavior, juveniles are considered one demographic group instead of two (female juvenile and male juvenile). We also considered adult females with dependent calves (mother-calf pairs) as one unit for disease transmission given the close contact between mother and calf. Further, due to our small sample size for PC of adult females without a dependent calf, we combined mother calf pairs and adult females into one demographic group to allow for a better balance of sample sizes across groups. For SB, we use the same demographic group definitions as PC to allow for reliable comparison (juvenile n=169; adult female and mother calf pair = 416; adult male = 25). Given high quality genetic data and high confidence birth observations for SB^72^, only focals with known demographic assignments were used, which resulted in 90% of contacts having known demographic assignments. Given that the SB population is demographically better-documented than PC, we also used SB data to support our demographic specific PC results. In Supplement Section 9, we used SB data to demonstrate that our demographic group definitions, which are constrained by the PC data limitations (i.e., juveniles are not sex differentiated; adult females with and without calves are grouped), are indeed appropriate. In Supplement Section 7, we showed that our results are robust to the uncertainty in PC demographic assignments.

### 4.2 Estimating transmission contact relevant to respiratory infection

Since DMV is a respiratory-transmitted pathogen, we focused on synchronized breathing interactions as our measure of contact. Past work in the field of network epidemiology demonstrates that average degree (the number of unique individuals an average focal individual has contact with), and degree heterogeneity (the variation among the population in the number of contacts per individual) are highly predictive of epidemic outcomes^28,73^. Additionally, we know that increased assortativity by trait (e.g., age or sex), or a tendency to have interactions with individuals of the same trait, can increase infection burden if the number of contacts are also structured by that trait^57^. Thus, to estimate disease outcomes among the PC bottlenose dolphins, we estimated synchrony degree, *k*_*d*_(*μ*) the number of unique individuals a focal animal of demographic group, *d*, has synchronous breathing interactions with over an average DMV infectious period, *μ*. We also estimated how the contacts of a focal individual are distributed among the demographic groups, *M*.

#### 2.2.1 Estimating demographic-specific synchrony degree

For contact to be a measure of the total number of individuals that an infected animal can transmit to while they are infected, it must be considered on the timescale of the infectious period of the pathogen. Since our focal follow data only measures synchronized breathing for short periods of time (average 25 minutes for PC and 119 for SB), it is not representative of total contact over a relevant infectious period for DMV (5-10 days^15^). To measure relevant synchrony degree, we must therefore extrapolate our measurements for longer time periods. It is challenging to extrapolate degree beyond periods of observation as there is fidelity in social bonding and interactions with the same individual are generally repeated. Based on past work, a power law relationship exists between degree and the number of interactions (while degree measures synchronous breathing events with unique individuals, interaction counts measure all synchronous breathing events) that spans behaviors and can be found across taxa^74^. So, we fit the following power law relationship to observed data on the number of synchronous breathing interactions that occur in *t* days (*n*_*d*_(*t*)), and synchrony degree per individual over *t* days (*k*_*d*_(*t*)) for each demographic group, *d*:

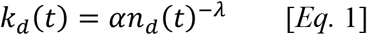

We identified the following parameters: *α* = *0*.*587* and *λ* = −*0*.*5209*.

Then, to empirically estimate the degree of an individual for a DMV infectious period *μ, k*_*d*_(*μ*), we must first estimate the number of synchrony interactions that individuals would have over *μ* days: *n*_*d*_(*μ*). To estimate *n*_*d*_(*μ*) we first decomposed the social processes required to be:

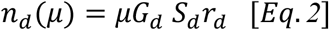

where *G*_*d*_ was the probability that individuals of demographic group *d* are observed in a group, *S*_*d*_ was the probability that individuals of demographic group *d* breathe synchronously while in a group, *r*_*d*_ was the typical rate of synchronized breathing for demographic group *d*, and *μ* was the infectious period of DMV. Next, using empirical data from each research site (PC and SB), we used a bootstrapping approach to estimate a distribution for *G*_*d*_ (based on survey data), and fit a series of generalized linear mixed models to estimate distributions for *S*_*d*_ (based on focal data), and *r*_*d*_(based on focal data). We then estimated *n*_*d*_(*μ*) by sampling from these fitted distributions for an infectious period, *με*[*5,10*] days for n=1000 individuals of each group *d*.

Using these estimates for *n*_*d*_(*μ*) and the power law fit from Eq. 1, we inferred *k*_*d*_(*μ*) and tested for statistically significant differences using a one-way ANOVA and Tukey’s test. Finally, we fit each *k*_*d*_(*μ*) mean and standard deviation to a negative binomial distribution as an estimate of the demographic-specific degree distribution *p*(*k*_*d*_(*μ*)), as this represents both the average degree and degree heterogeneity for each group *d*. Full details on 2.2.1 methods can be found in Supplemental section 3.

### Validation

To demonstrate that our estimate of degree by demographic group over an infectious period *k*_*d*_(*μ*) is supported by data, we compared our *k*_*d*_(*μ*) to our observations of degree during an average focal follow period^74^ show that the (power law) relationship between degree and the number of interactions is determined by the nature of social behaviors (affiliative versus antagonistic) and is relatively consistent. Given that our study focuses on a particular (affiliative) behavior, we assumed that the relationship between number of interactions and degree is stable for a given demographic group, and that our estimated degree for an infectious period should reflect similar patterns to the observed degree (PC n=99 follows, SB n=610 follows) over an average focal follow period (for further details, see Supplement Section 3.5). To control for the length of the follow, and variation across years and individuals, we ran a generalized linear mixed model with observed degree as the response variable and a categorical predictor variable for demographic groups with the length of the follow as a fixed effect, and random effects for year and dolphin identity.

#### 4.2.2 Estimating demographic mixing in synchronous breathing

To evaluate the degree of mixing among demographic groups, we measured demographic assortativity (the degree to which individuals interact with individuals of their own demographic group) by creating a mixing matrix based on focal follow data (PC: n=99, SB: n=610). Each element *m*_*gh*_ in *M* represented the proportion of contacts for a focal individual of demographic group *g* that are of demographic group *h*, across all follows. We standardized each *m*_*gh*_ term to control for differences in the number of individuals within each demographic group and their estimated *k*_*d*_(*μ*), and then used a bootstrapping approach to estimate confidence intervals for each term. Finally, we generated a matrix *E* to represent our expectation for demographic mixing assuming it occurred at random (for details, see Supplement Section 4). A comparison of *M* to *E* gives us an understanding of which demographic pair is mixing more than expected. We also calculated the assortativity coefficients for *M* according to Newman’s definition ^75^. To test if dolphins are more assortative than expected at random, we transformed the assortativity coefficients into z scores and tested for statistical significance.

### 4.3 Network Generation

Informed by our estimates of demographic-specific synchrony contact structure, we constructed contact network models that represent synchronous breathing contacts unique to each study site. A network was characterized by nodes representing individual dolphins (or mother-calf pairs), and edges representing the presence of synchronized breathing contact between a pair of individuals. Based on the demographic-specific synchrony degree distributions (*p*(*k*_*d*_(*μ*))), synchrony demographic mixing matrix (*M*), and demographic distribution of the population (specified in Supplement Section 2), we developed a network generating algorithm to create synthetic networks representing the contact structure of bottlenose dolphins. Networks of any size can be created with our algorithm, but we specified networks of 2000 nodes (individuals) in order to reflect reasonable estimates of the number of individuals sighted in the PC and SB ^51^. The synthetic networks were random in all other properties besides those specified (degree distribution by demographic group, and mixing patterns between and among groups). This feature is critical as it allows us to study the implications of the specified structure in a systematic way. We generated 25 networks to capture variability in contact dynamics.

### 4.4 Modeling disease transmission in bottlenose dolphins

To model the spread of DMV across our generated network models, we performed numerical simulations of a susceptible-infected-recovered percolation model^75^. For each network, we simulated 100 disease outbreaks where infection started in each demographic group, and classified large-scale epidemics as those where at least 10% of the network is infected. The average R0 for morbilliviruses across marine mammals is 1.9 ^15,41,76^. We evaluated disease outcomes by calculating the 1) epidemic size (proportion of network infected in an epidemic), and 2) demographic risk (proportion of each demographic group infected in an epidemic) across all simulations regardless of the demographic group of the originally infected node. We tested for statistically significant differences between age and sex class infection risks using one-tailed t-tests. We showed that the variability of epidemic outcomes is not different within one network, compared to across all 25 generated networks to demonstrate that we generated enough networks to capture potential contact variability (see fig. S15).

#### 4.4.1 Validation from empirical disease data

To demonstrate that our model results are supported by observed disease data, we obtained mortality data available for bottlenose dolphins that washed ashore or were found floating, defined as “stranded” from 2010-2015 from the US National Marine Fisheries Service. The DMV epidemic spanned 21 months (1-Jul-13 to 1-Mar-15). Because not all recorded cases of strandings during this time period (n=1,263 strandings) would be due to DMV, we defined excess mortality by removing typical “background” mortality for each demographic group (such as from natural mortality or human interactions) ^15,77^. The resulting data therefore represented the number of recovered mortalities that occurred due to DMV along the Atlantic coast, which we assumed is proportional to the number of individuals infected during the epidemic (n=504 strandings). To make these data comparable to our model results, we standardized the number of deaths for each demographic class to be an estimated proportion of recovered epidemic mortality for that age or sex class for Atlantic dolphins (See Supplement Section 5 for more details this methodology). We tested for statistically significant differences between age and sex class strandings using a one-tailed t-test. These methods assume that the effort to collect dolphin stranding data did not change from before the outbreak to during the outbreak. While this assumption may not be true, we did not find any evidence to suggest that effort would affect the demographic trends of strandings so we suggest this assumption is valid for our purposes.

#### 4.4.2 Demographic-specific disease introduction

We were also interested in how epidemic outcomes change based on where in the network the pathogen is introduced. While DMV is not a reverse zoonotic pathogen, there are instances in which wild dolphins have direct contact with humans that could allow for emergence of other respiratory transmitted pathogens in dolphins, such as through human associated foraging and in swim with dolphin programs ^78–80^. There has also been observed variation in how frequently different demographics partake in human interactions. For example, studies have found that juvenile bottlenose dolphins are more likely to come into close contact with humans participating in wild swim with dolphin tourism ^81^. Further, adult male cetaceans in certain populations are more likely to interact with humans by depredating recreational fishing lines ^82^ and food provisioning ^83^. Thus, we assessed the impact of disease introductions in particular demographic groups on disease risk. Because disease introductions are known to affect the likelihood of an epidemic occurring, or epidemic probability, rather than epidemic size, we considered a product of epidemic probability and demographic risk and tested for differences between scenarios using a one-way ANOVA and Tukey’s test.

#### 4.4.3 Structural analysis

To examine which structural feature of the contact network is most predictive of demographic specific risk, we defined two null networks: one in which demographic-specific degree distributions were random and homogeneous across demographic groups but demographic mixing was still empirically-informed; and another in which demographic mixing was random, but demographic-specific degree distributions were retained as empirically-informed. We compared disease consequences of our empirically-informed networks against these two null models (see Supplement Section 8 for details) using a one-way ANOVA and Tukey’s test.

## Acknowledgements

We thank the Potomac-Chesapeake Dolphin Project field assistants for 2018-2022 without whom data collection would not have been possible: 2022-Marley Dooling, Meng-Chun Grace Chung; 2021-Sarah Theisen, Amelia Smith, Katie Knotek, Sophie Hanson; 2020-Milan Dolezal; 2019-Trevyn Toone, Molly Albright, Haley Land-Miller, Casey Marker, Kelsey O’Donnell; 2018-Jessica Wang, Jazmin Garcia. We also thank our collaborators and field assistants on the Shark Bay Dolphin Research Project and the West Australian Department of Biodiversity, Conservation, and Attractions. We thank the members of the Greater Atlantic and Southeast Regional Marine Mammal Stranding Networks for the collection of the stranding data used, and the National Marine Fisheries Service for providing this data.

All PC and Shark Bay data were collected under GU Permit #s IACUC-13-069, 07-041, 10-023; 2016-1235. PC data were collected under NMFS permit nos 19403 and 23782. Shark Bay data were collected under permits from the West Australian Department of Biodiversity, Conservation and Attractions: SF-009876, SF-010347, SF-008076, SF009311, SF007457, FO25000102-6.

## Author Contributions

Conceptualization: MAC, SB

Data Collection: MAC, A-MJ, VF, EMP, EK, MMW, MM, CK, JM

Data Contribution: A-MJ, VF, JM, SW, SB Analysis Tool Contribution: MAC, SB Analysis: MAC, SB

Supervision: JM, SB Writing—original draft: MAC

Writing—review & editing: MAC, A-MJ, VF, EMP, MMW, MM, CK, JM, SB

## Data and Materials Availability

All data needed to evaluate the conclusions in the paper are present in the paper, the Supplementary Materials, and at https://zenodo.org/doi/10.5281/zenodo.11034494

## References

1. Gulland, F. M. D. et al. A review of climate change effects on marine mammals in United States waters: Past predictions, observed impacts, current research and conservation imperatives. Clim. Change Ecol. 3, 100054 (2022).

2. Bongaarts, J. IPBES, 2019. Summary for policymakers of the global assessment report on biodiversity and ecosystem services of the Intergovernmental Science-Policy Platform on Biodiversity and Ecosystem Services. Popul. Dev. Rev. 45, 680–681 (2019).

3. Cohen, R. E. et al. Marine host-pathogen dynamics: Influences of global climate change. Oceanogr. Soc. 31, 182–193 (2018).

4. Cook, T., Folli, M., Klinck, J., Ford, S. & Miller, J. The relationship between increasing sea surface temperature and the northward spread of Perkinsus marinus (Dermo) disease epizootics in oysters. Estuar. Coast. Shelf Sci. 46, 587–597 (1998).

5. Conrad, P. A. et al. Transmission of Toxoplasma: Clues from the study of sea otters as sentinels of Toxoplasma gondii flow into the marine environment. Int. J. Parasitol. 35, 1155– 1168 (2005).

6. Schaefer, A. M. et al. Serological evidence of exposure to selected viral, bacterial, and protozoal pathogens in free-ranging Atlantic bottlenose dolphins (Tursiops truncatus) from the Indian River Lagoon, Florida, and Charleston, South Carolina. Aquat. Mamm. 35, 163– 170 (2009).

7. Bossart, G. D. Marine Mammals as sentinel species for ocean and human health. Oceanography 19, 134–137 (2006).

8. Heithaus, M. R., Frid, A., Wirsing, A. J. & Worm, B. Predicting ecological consequences of marine top predator declines. Trends Ecol. Evol. 23, 202–210 (2008).

9. Moore, S. Marine mammals as ecosystem sentinels. J. Mammology 89, 534–540 (2008).

10. Gulland, F. M. D. & Hall, A. J. Is marine mammal health deteriorating? Trends in the global reporting of marine mammal disease. EcoHealth 4, 135–150 (2007).

11. Van Bressem, M. F. et al. Cetacean morbillivirus: current knowledge and future directions. Viruses 6, 5145–5181 (2014).

12. Silk, M. J. et al. Contact networks structured by sex underpin sex-specific epidemiology of infection. Ecol. Lett. 21, 309–318 (2018).

13. Hayes, S. A. et al. U.S. Atlantic and Gulf of Mexico Marine Mammal Stock Assessments 2022. NOAA Tech Memo NMFS NE 304 (2023).

14. NOAA Fisheries. Depleted Designation for Western North Atlantic Coastal Migratory Stock of Bottlenose Dolphins. https://www.fisheries.noaa.gov/action/depleted-designation-western-north-atlantic-coastal-migratory-stock-bottlenose-dolphins (1993).

15. Morris, S. E. et al. Partially observed epidemics in wildlife hosts: modelling an outbreak of dolphin morbillivirus in the northwestern Atlantic, June 2013–2014. J. R. Soc. Interface 12, (2015).

16. NOAA. Active and Closed Unusual Mortality Events. NOAA Fisheries https://www.fisheries.noaa.gov/national/marine-life-distress/active-and-closed-unusual-mortality-events (2020).

17. Keck, N. et al. Resurgence of Morbillivirus infection in Mediterranean dolphins off the French coast. Vet. Rec. 166, 654–655 (2010).

18. Hosseini, P. R., Dhondt, A. A. & Dobson, A. Seasonality and wildlife disease: How seasonal birth, aggregation and variation in immunity affect the dynamics of Mycoplasma gallisepticum in house finches. Proc. R. Soc. B Biol. Sci. 271, 2569–2577 (2004).

19. Collier, M. A., Mann, J., Ali, S. & Bansal, S. Impacts of Human Disturbance in Marine Mammals: Do Behavioral Changes Translate to Disease Consequences?. in Marine Mammals: the Evolving Human Factor. Ethology and Behavioral Ecology of Marine Mammals (eds. Wursig, B. & Di Sciara, G. N.) 277–305 (Springer, 2022). doi:10.1007/978-3-030-98100-6_9.

20. Bansal, S., Grenfell, B. T. & Meyers, L. A. When individual behaviour matters: homogeneous and network models in epidemiology. J. R. Soc. Interface 4, 879–891 (2007).

21. Leu, S. T. et al. Sex, synchrony, and skin contact: integrating multiple behaviors to assess pathogen transmission risk. Behav. Ecol. 31, 651–660 (2020).

22. Connor, R. C., Wells, R., Mann, J. & Read, A. Social relationships in a fission-fusion society. in Cetacean Societies: Field Studies of Dolphins and Whales (eds. Mann, J., Connor, R., Tyack, P. & Whitehead, H.) 91–126 (Chicago: The University of Chicago Press., 2000).

23. Galezo, A. A., Krzyszczyk, E. & Mann, J. Sexual segregation in Indo-Pacific bottlenose dolphins is driven by female avoidance of males. Behav. Ecol. 29, 377–386 (2018).

24. Tsai, Y.-J. J. & Mann, J. Dispersal, philopatry, and the role of fission-fusion dynamics in bottlenose dolphins. Mar. Mammal Sci. 29, 261–279 (2013).

25. Connor, R. C., Smolker, R. & Bejder, L. Synchrony, social behaviour and alliance affiliation in Indian Ocean bottlenose dolphins, Tursiops aduncus. Anim. Behav. 72, 1371– 1378 (2006).

26. Foroughirad, V. et al. Small effects of family size on sociality despite strong kin preferences in female bottlenose dolphins. Anim. Behav. 195, 53–66 (2023).

27. Krzyszczyk, E., Patterson, E. M., Stanton, M. A. & Mann, J. The transition to independence: sex differences in social and behavioural development of wild bottlenose dolphins. Anim. Behav. 129, 43–59 (2017).

28. Collier, M. A., Albery, G. F., Mcdonald, G. C. & Bansal, S. Pathogen transmission modes determine contact network structure, altering other pathogen characteristics. Proc. R. Soc. B Biol. Sci. 289, 0221389 (2022).

29. Brager, S. Diurnal and seasonal behavior patterns of bottlenose dolphins (Tursiops truncatus). Mar. Mammal Sci. 9, 434–438 (1993).

30. Hanson, M. T. & Defran, R. H. The behaviour and feeding ecology of the Pacific coast bottlenose dolphin, Tursiops truncatus. Aquat. Mamm. 19, 127–142 (1993).

31. Mann, J. Behavioral sampling methods for cetaceans: a review and critique. Mar. Mammal Sci. 15, 102–122 (1999).

32. Moller, L. M. & Harcourt, R. G. Social dynamics and activity patterns of bottlenose dolphins, Tursiops truncatus, in Jarvis Bay, S.E. Australia. Proc. Linneaen Soc. New South Wales 120, 181–189 (1998).

33. Sakai, M., Morisaka, T., Kogi, K., Hishii, T. & Kohshima, S. Fine-scale analysis of synchronous breathing in wild Indo-Pacific bottlenose dolphins (Tursiops aduncus). Behav. Process. 83, 48–53 (2009).

34. Hastie, G. D., Wilson, B., Tufft, L. H. & Thompson, P. M. Bottlenose dolphins increase breathing synchrony in response to boat traffic. Mar. Mammal Sci. 19, 74–84 (2003).

35. Simila, T. Sonar observations of killer whales (Orcinus Orca) feeding on herring schools. Aquat. Mamm. 23, 119–126 (1997).

36. White, L. A., Forester, J. D. & Craft, M. E. Using contact networks to explore mechanisms of parasite transmission in wildlife. Biol. Rev. 92, 389–409 (2017).

37. Ben-Horin, T. et al. Modelling marine diseases. in Marine Disease Ecology (eds. Behringer, D. C., Silliman, B. R. & Lafferty, K. D.) 233–256 (Oxford University Press, 2020). doi:10.1093/oso/9780198821632.003.0012.

38. Keeling, M. J. & Rohani, P. Modeling Infectious Disease. (Princeton University Press, Princeton, 2008).

39. Baker, J. D. et al. Modeling a morbillivirus outbreak in Hawaiian Monk seals (Neomonachus schauinslandi) to aid in the design of mitigation programs. J. Wildl. Dis. 53, 736–748 (2017).

40. Bodewes, R. et al. Prevalence of phocine distemper virus specific antibodies: bracing for the next seal epizootic in north-western Europe. Emerg. Microbes Infect. 2, e3 (2013).

41. De Koeijer, A., Diekmann, O. & Reijnders, P. Modelling the spread of phocine distemper virus among harbour seals. Bull. Math. Biol. 60, 585–596 (1998).

42. Harris, C. M., Travis, J. M. J. & Harwood, J. Evaluating the influence of epidemiological parameters and host ecology on the spread of phocine distemper virus through populations of harbour seals. PLOS ONE 3, e2710 (2008).

43. Heide-Jorgensen, M.-P. & Harkonen, T. Epizootiology of the seal disease in the eastern North Sea. J. Appl. Ecol. 29, 99–107 (1992).

44. Swinton, J., Harwood, J., Grenfell, B. T. & Gilligan, C. A. Persistence thresholds for phocine distemper virus infection in harbour seal Phoca vitulina metapopulations. J. Anim. Ecol. 67, 54–68 (1998).

45. Costa, A. P. B., Mcfee, W., Wilcox, L. A., Archer, F. I. & Rosel, P. E. The common bottlenose dolphin (Tursiops truncatus) ecotypes of the western North Atlantic revisited: an integrative taxonomic investigation supports the presence of distinct species. Zool. J. Linn. Soc. 196, 1608–1636 (2022).

46. Waring, G. T., Josephson, E., Maze-Foley, K. & Rosel, P. E. US Atlantic and Gulf of Mexico Marine Mammal Stock Assessments 2015. NOAA Tech Memo NMFS NE 238 501 (2016).

47. Kirk, C. M., Amstrup, S., Swor, R., Holcomb, D. & O’Hara, T. M. Morbillivirus and Toxoplasma exposure and association with hematological parameters for southern Beaufort Sea polar bears: potential response to infectious agents in a sentinel species. EcoHealth 7, 321–331 (2010).

48. Pomeroy, P. P. et al. Morbillivirus neutralising antibodies in Scottish grey seals Halichoerus grypus: assessing the effects of the 1988 and 2002 PDV epizootics. Mar. Ecol. Prog. Ser. 287, 241–250 (2005).

49. Tango, P. J. & Batiuk, R. A. Chesapeake Bay recovery and factors affecting trends: Long-term monitoring, indicators, and insights. Reg. Stud. Mar. Sci. 4, 12–20 (2016).

50. Lipscomb, T. P., Schulman, F. Y., Moffett, D. & Kennedy, S. Morbilliviral disease in Atlantic bottlenose dolphins (Tursiops truncatus) from the 1987-1988 epizootic. J. Wildl. Dis. 30, 567–571 (1994).

51. Preen, A. R., Marsh, H., Lawler, I. R., Prince, R. I. T. & Shepherd, R. Distribution and abundance of dugongs, turtles, dolphins and other megafauna in Shark Bay, Ningaloo Reef and Exmouth Gulf, western Australia. Wildl. Res. 24, 185–208 (1997).

52. Barco, S. G., Swingle, W. M., Mlellan, W. A., Harris, R. N. & Pabst, D. A. Local abundance and distribution of bottlenose dolphins (Tursiops truncatus) in the nearshore waters of Virginia Beach, Virginia. Mar. Mammal Sci. 15, 394–408 (1999).

53. Engelhaupt, A., Jefferson, T. A., Aschettino, J. M. & Bell, J. T. Distribution, abundance and sighting patterns of multiple stocks of bottlenose dolphins (Tursiops truncatus) in coastal Virginia waters. J Cetacean Res Manage 23, 109–125 (2022).

54. Rodriguez, L. K., Fandel, A. D., Colbert, B. R., Testa, J. C. & Bailey, H. Spatial and temporal variation in the occurrence of bottlenose dolphins in the Chesapeake Bay, USA, using citizen science sighting data. PLOS ONE 16, e0251637 (2021).

55. Mann, J. & Sargeant, B. Like mother, like calf: the ontogeny of foraging traditions in wild Indian Ocean bottlenose dolphins (Tursiops sp.). in The Biology of Traditions: Models and Evidence (eds. Fragaszy, D. M. & Perry, S.) 236–266 (Cambridge University Press, 2003).

56. Karniski, C. et al. A comparison of survey and focal follow methods for estimating individual activity budgets of cetaceans. Mar. Mammal Sci. 31, 839–852 (2015).

57. Wallinga, J., Teunis, P. & Kretzschmar, M. Using data on social contacts to estimate age-specific transmission parameters for respiratory-spread infectious agents. Am. J. Epidemiol. 164, 936–944 (2006).

58. Miller, L. J., Solangi, M. & Kuczaj, S. A. Immediate response of Atlantic bottlenose dolphins to high-speed personal watercraft in the Mississippi Sound. Mar. Biol. Assoc. U. K. J. Mar. Biol. Assoc. U. K. 88, 1139–1143 (2008).

59. Evans, T., Krzyszczyk, E., Frère, C. & Mann, J. Lifetime stability of social traits in bottlenose dolphins. Commun. Biol. 4, (2021).

60. Smolker, R. A., Richards, A. F., Connor, R. C. & Pepper, J. W. Sex differences in patterns of association among Indian Ocean bottlenose dolphins. Behaviour 123, 38–69 (1992).

61. Ferrari, N., Cattadori, I. M., Nespereira, J., Rizzoli, A. & Hudson, P. J. The role of host sex in parasite dynamics: field experiments on the yellow-necked mouse Apodemus flavicollis. Ecol. Lett. 7, 88–94 (2004).

62. Godfrey, S. S. Networks and the ecology of parasite transmission: A framework for wildlife parasitology. Int. J. Parasitol. Parasites Wildl. 2, 235–245 (2013).

63. Frere, C. H. et al. Social and genetic interactions drive fitness variation in a free-living dolphin population. Biol. Sci. 107, 19949–19954 (2010).

64. Gerber, L. et al. Social integration influences fitness in allied male dolphins. Curr. Biol. 32, 1664–1669 (2022).

65. Wiszniewski, J., Corrigan, S., Beheregaray, L. B. & Möller, L. M. Male reproductive success increases with alliance size in Indo-Pacific bottlenose dolphins (Tursiops aduncus). J. Anim. Ecol. 81, 423–431 (2012).

66. Galezo, A. A., Foroughirad, V., Krzyszczyk, E., Frere, C. H. & Mann, J. Juvenile social dynamics reflect adult reproductive strategies in bottlenose dolphins. Behav. Ecol. 31, 1159–1171 (2020).

67. Leguia, M. et al. Highly pathogenic avian influenza A (H5N1) in marine mammals and seabirds in Peru. Nat. Commun. 14, (2023).

68. Thorsson, E. et al. Highly pathogenic avian influenza A(H5N1) virus in a harbor porpoise, Sweden. Emerg. Infect. Dis. 29, 852–855 (2023).

69. Nelson, T. M. et al. Detecting respiratory bacterial communities of wild dolphins: implications for animal health. Mar. Ecol. Prog. Ser. 622, 203–217 (2019).

70. Vaughan, K. et al. A DNA vaccine against dolphin morbillivirus is immunogenic in bottlenose dolphins. Vet. Immunol. Immunopathol. 120, 260–266 (2007).

71. Hammond, P., Mizroch, S. & Donovan, G. Individual Recognition of Cetaceans: Use of Photo-Identification and Other Techniques to Estimate Population Parameters. In: Report of the International Whaling Comission). https://www.semanticscholar.org/paper/Individual-recognition-of-cetaceans%3A-use-of-and-to-Hammond-Mizroch/691e0e729d5e8d2665b34af427e38687a1ad4c2a (1990).

72. McEntee, M. H. F., Foroughirad, V., Krzyszczyk, E. & Mann, J. Sex bias in mortality risk changes over the lifespan of bottlenose dolphins. Proc. R. Soc. B Biol. Sci. 290, 20230675 (2023).

73. Sah, P., Mann, J. & Bansal, S. Disease implications of animal social network structure: A synthesis across social systems. J. Anim. Ecol. 1–13 (2018) doi:10.1111/1365-2656.12786.

74. Colman, E., Colizza, V., Hanks, E. M., Hughes, D. P. & Bansal, S. Social fluidity mobilizes contagion in human and animal populations. eLife 10, e62177 (2021).

75. Newman, M. E. J. Spread of epidemic disease on networks. Phys. Rev. E - Stat. Phys. Plasmas Fluids Relat. Interdiscip. Top. 66, 016128 (2002).

76. Lonergan, M. et al. Comparison of the 1988 and 2002 phocine distemper epizootics in British harbour seal Phoca vitulina populations. Dis. Aquat. Organ. 88, 183–188 (2010).

77. Farr, W. Vital Statistics: A Memorial Volume of Selections From the Reports and Writings of William Farr London. (Offices of the Sanitary Institute, 1885).

78. Cunningham-Smith, P., Colbert, D. E., Wells, R. S. & Speakman, T. Evaluation of human interactions with a provisioned wild bottlenose dolphin (Tursiops truncatus) near Sarasota Bay, Florida, and efforts to curtail the interactions. Aquat. Mamm. 32, 346–356 (2006).

79. Hazelkorn, R. A., Schulte, B. A. & Cox, T. M. Persistent effects of begging on common bottlenose dolphin (Tursiops truncatus) behavior in an estuarine population. Aquat. Mamm. 42, 531–541 (2016).

80. Peters, K. J., Parra, G. J., Skuza, P. P. & Möller, L. M. First insights into the effects of swim-with-dolphin tourism on the behavior, response, and group structure of southern Australian bottlenose dolphins. Mar. Mammal Sci. 29, 484–497 (2013).

81. Constantine, R. Increased avoidance of swimmers by wild bottlenose dolphins (Tursiops truncatus) due to long-term exposure to swim-with-dolphin tourism. Mar. Mammal Sci. 17, 689–702 (2001).

82. Powell, J. R. & Wells, R. S. Recreational fishing depredation and associated behaviors involving common bottlenose dolphins (Tursiops truncatus) in Sarasota Bay, Florida. Mar. Mammal Sci. 27, 111–129 (2011).

83. Pinto De Sá Alves, L. C., Andriolo, A., Orams, M. B. & De Freitas Azevedo, A. Resource defence and dominance hierarchy in the boto (Inia geoffrensis) during a provisioning program. Acta Ethologica 16, 9–19 (2013).

84. Mann, J. & Smuts, B. Behavioral developments in wild bottlenose dolphin newborns (Tursiops sp.). Behaviour 136, 529–566 (1999).

85. Connor, R. C., Wells, R. S., Mann, J. & Read, A. J. The bottlenose dolphin. Cetacean Soc. 91–25 (2000).

86. Tolley, K. A. et al. Sexual dimorphism in wild bottlenose dolphins (Tursiops truncatus) from Sarasota, Florida. J. Mammal. 76, 1190–1198 (1995).

87. Read, A. J., Wells, R. S., Hohn, A. A. & Scott, M. D. Patterns of growth in wild bottlenose dolphins, Tursiops truncatus. J. Zool. 231, 107–123 (1993).

88. Mann, J. Establishing Trust: Sociosexual behaviour and the development of male-male bonds among Indian Ocean bottlenose dolphin calves. in Animals: An Evolutionary Perspective (eds. Vasey, P. & Sommer, P.) 107–130 (Cambridge University Press, 2006).

89. Schroeder, J. P. Breeding Bottlenose Dolphins in Captivity. in Handbook of Marine Mammals (eds. Ridgeway, S. H. & Harrison, R. J.) 435–446 (Academic Press, 1999).

90. Hohn, A. A. Age determination and age related factors in the teeth of western north Atlantic bottlenose dolphins. Sci. Rep. Whales Res. Institue Tokyo 32, 39–66 (1980).

91. Mead, J. G. & Potter, C. W. Natural history of bottlenose dolphins along the central Atlantic coast of the United States. in The bottlenose dolphin (eds. Leatherwood, S. & Reeves, R. R.) 165–198 (Academic Press, San Diego, CA, 1990).

92. Cockcroft, V. G. & Ross, G. J. B. Age, growth, and reproduction of bottlenose dolphins Tursiops truncatus from the east coast of southern Africa. Fish. Bull. 88, 289–302 (1990).

