## Supplementary Information for "Breathing in sync: how a social behavior structures respiratory epidemic risk in bottlenose dolphins"

### Section 1: Assigning focal individuals and contacts to a demographic group for the Potomac-Chesapeake Dolphin Project (PC)

We considered each individual as a member of one of three demographic groups: adult male, adult female (including mother-calf pairs, which are treated as one unit) and juvenile (table S1). Our age and sex class rules (see Appendix 1 below) allowed us to assign age and sex class to an individual with both high and low confidences. However, if an individual did not meet the requirements for even a low confidence age or sex class assignment, it was considered to be of 'unknown' age or sex class.

Assigning individuals with unknown age and sex classes:

Focal individuals assigned as an unknown age class were dropped from this analysis. Focal individuals classified as adults but of unknown sex were also dropped from this analysis. However, we still included individuals of unknown age or sex class who had contact with focals when developing our demographic mixing matrices. Therefore, we generated a 'confirmed' demographic mixing matrix using only focals where all contacts had been assigned to a demographic group with high or low confidence. Then, for focals with unknown contacts, we assigned each unknown contact to one of our three demographic groups based on: the focal's demographic group and the likelihood that the focal interacts with other demographic groups based on the values in the confirmed mixing matrix. The resulting demographic assignment of these unknowns was designated as low confidence. We used these random assignments to generate our full demographic mixing matrix, and performed a sensitivity analysis to assess the robustness of our results to incorrect demographic group assignments for all low confidence individuals (see Section 7 for results).

### Section 2: Demographic Distributions

Using life history data from the SB, we calculated the distribution of our demographic groups across the SB population as an average of 42% adult male, 44% adult female and mother calf pairs and 14% juveniles each year. We used this demographic distribution to generate our synthetic contact networks for the SB. We also assumed that the distribution of demographic groups for the PC was the same, and generated our synthetic networks based on this breakdown.

### Section 3: Detailed methodology and results for estimating $k_d(\mu)$

Our focal follow data only examined synchrony behavior over a two hour period on average, which is not representative of contact over a relevant infectious period for DMV. Since bottlenose dolphins live in gregarious fission-fusion societies [23,24], we would expect to see their synchrony degree increase over time. However, because synchronized breathing is an affiliative behavior most commonly seen between closely bonded individuals, we would not expect this relationship to be truly linear. Established research [74] suggests that the rate of social mixing in a population can be described by a single mathematical parameter known as social fluidity where large values correspond to more gregarious behavior, and small values signify the existence of persistent bonds between individuals (see fig. S2). Therefore, as assumed in Colman et al., we assumed that that the synchrony degree over infectious period γ ($k(\mu)$) could be measured using a power law relationship with the number of synchronized breathing interactions that occur over μ, $n(\mu)$:

$$k(\mu) \sim \alpha\, n(\mu)^{-\lambda}$$

Where α and λ were parameters representing bottlenose dolphin social fluidity. To estimate α and λ, we used the R package movr to fit a power law function to our observed focal follow data. We first examined the social fluidity for both the PC and SB separately but found very little difference between the two relationships when the number of synchronized breaths is well represented for both study sites (fig. S3). Since the resulting rate only diverged where we lack data for the PC (due to shorter follow lengths), we developed one curve for both populations, and estimated social fluidity parameters as $\alpha = 0.587;\ \lambda = \ -0.5209$ (fig. S4). We then assumed that the average degree over an infectious period for each demographic group $d$ $k_d(\mu)$ could be estimated as:

$$k_d(\mu) = 0.587 n_d(\mu)^{0.5209}$$

We then assumed that $n_d(\mu)$ would depend on 1) the probability that demographic group $d$ is observed in a group $G_d$, 2) the probability that demographic group $d$ breathes synchronously while in a group $S_d$, and 3) the rate of breathing synchrony $r_d$ for demographic group $d$. Therefore given infectious period γ, we estimated:

$$n_d(\mu) = \mu G_d S_d r_d$$

<u>3.1 Estimating $G_d$</u>

We estimated the probability of being in a group for each demographic $G_d$ using survey data. For the SB, we used all surveys conducted during the same time period as our focal follows (2010-2018) for which age and sex was known for all individuals (n = 5,103). We determined the probability that each demographic was alone $A_d$ to be the number of surveys on one individual (or one mother/calf pair) in the demographic group divided by the total number of surveys that include that demographic group. We then calculated $G_d = 1 - A_d$.

For the PC, we did not have the ability to assign a demographic group to every individual in every survey. Therefore, we assumed that the demographic trends of $G_d$ in PC were proportional to that of SB, but the magnitude of $G_d$ may have differed between the two study sites. Therefore, we estimated:

$$A_{PC/d} = \frac{\frac{a_{PC} * a_{SB/d}}{a_{SB}}}{\frac{n_{PC} * n_{SB/d}}{n_{SB}}}$$

Where $a$ was the number of surveys on lone individuals (or mother calf pairs) and $n$ was the total number of surveys. Sample sizes can be found in table S2.

For both the PC and the SB we used a bootstrapping approach to generate confidence intervals for $G_d$.

Results:
For the SB we estimate $G_d$ as:

$$G_{adult\ male} = 0.76\ [0.73 - 0.79]$$

$$G_{adult\ female} = 0.63\ [0.61 - 0.66]$$

$$G_{juvenile} = 0.77\ [0.73 - 0.80]$$

For the PC we estimate $G_d$ as:

$$G_{adult\ male} = 0.93\ [0.89 - 0.95]$$

$$G_{adult\ female} = 0.90\ [0.84 - 0.93]$$

$$G_{juvenile} = 0.93\ [0.90 - 0.96]$$

3.2 Estimating $S_d$

We estimated the probability that an individual will sync if it's in a group $S_d$ using focal follow data. We used a generalized linear mixed model with a binary response variable for whether a sync occurred during a follow. We used a categorical predictor variable for demographic group, a continuous effect for the length of the follow to control for increased likelihood of syncing over time, and random effects for the year the follow was conducted and the ID of the dolphin to control for variability in data collected across years, and individual behavior of the dolphin respectively. We then chose 1000 random model estimates for each demographic grouping to generate a distribution for each $S_d$.

Results:
In the PC, we find that adult males are more likely to sync when they are in a group compared to adult females. Juveniles are also more likely to sync than adult females on average, but the wide variation in $S_d$ makes this result not significant (fig. S5). We estimate $S_d$ for the PC as:

$S_{adult\ male} = 0.87\ [0.63 - 0.96]$

$S_{adult\ female} = 0.63\ [0.34 - 0.85]$

$S_{juvenile} = 0.88\ [0.42 - 0.97]$

In SB, we find that juveniles are more likely to sync than adult females when they are in a group. On average, adult males show a non-significant tendency to sync more than adult females, with wide variation in $S_d$ (fig. S6). We estimate $S_d$ for the SB as:

$S_{adult\ male} = 0.51\ [0.29 - 0.72]$

$S_{adult\ female} = 0.46\ [0.38 - 0.54]$

$S_{juvenile} = 0.69\ [0.58 - 0.78]$

3.3 Estimating $r_d$

We estimated the rate of synchrony for each demographic $r_d$ using focal follow data. We dropped any follows where a sync did not occur. We used a generalized linear mixed model with a continuous response variable for the number of syncs that occurred during a follow. We used a categorical predictor variable for demographic group and a continuous predictor for the length of the follow. We included random effects for the year the follow was conducted and the ID of the dolphin to control for variability in data collected across years, and individual behavior of the dolphin respectively. As the model estimates the number of syncs that will occur for each demographic over the average follow length in our dataset (24.6 minutes for PC and 119.1 minutes for SB) we calculated $r_d$ by choosing 1000 random model estimates for each demographic group and divided it by the average follow length to generate a distribution of each $r_d$.

Results

In the PC, we find that juveniles have the highest $r_d$, followed by adult males and adult females (fig. S7). We estimate $r_d$ for the PC as:

$$r_{adult\ male} = 0.24\ [0.19,\ 0.29]\ syncs\ per\ minute$$

$$r_{adult\ female} = 0.19\ [0.15 - 0.24]\ syncs\ per\ minute$$

$$r_{juvenile} = 0.32\ [0.24 - 0.44]\ syncs\ per\ minute$$

In the SB, we find that adult males have the highest $r_d$, followed by juveniles and adult females (fig. S8). We estimate $r_d$ for the SB as:

$$r_{adult\ male} = 0.09\ [0.05 - 0.17]\ syncs\ per\ minute$$

$$r_{adult\ female} = 0.04\ [0.03 - 0.04]\ syncs\ per\ minute$$

$$r_{juvenile} = 0.05\ [0.04 - 0.06]\ syncs\ per\ minute$$

3.5 Validating our estimate of $k_d(\mu)$ with observed synchrony degree

We support our $k_d(\mu)$ estimates by demonstrating the observed synchrony degree for each demographic $g$ over the average focal follow length. We used a generalized linear mixed model, where our response variable was the synchrony degree for each follow, with a categorical fixed effect for demographic group, and a continuous fixed effect for the length of the follow. We also included random effects for the year of the follow, and the ID of the dolphin to control for variability in data collected across years, and individual behavior of the dolphin respectively. We compared the model estimates for observed synchrony degree to our estimated $k_d(\mu)$ to make sure that the demographic specific patterns observed from our theoretical $k_d(\mu)$ are also represented empirically. Results for PC are found in the main text Fig. 2. Results for SB are found in fig. S10.

**Section 4: Detailed methods and results for estimating demographic assortative mixing over a DMV infectious period**

4.1 Calculating $M$ and $E$

In the mixing matrix $M$, each row represented the demographic group of the focal individuals, and each column represented the demographic group of their contacts. Each element $m_{gh}$ in $M$ represented the proportion of contacts for a focal individual of demographic group $g$ that are of demographic group $h$, across all follows. We standardized each $m_{gh}$ to control for population differences in the number of individuals within each demographic group and their estimated $k_g(\mu)$: $(m_{gh}(\mu))$. We estimated $m_{gh}(\mu)$ as:

$$m_{gh}(\mu) = m_{gh} \frac{k_g(\mu)p_g}{k(\mu)}$$

Where $k_g(\mu)$ was the average estimated degree for demographic group $g$ for infectious period $D$, $p_g$ was the proportion of the total population that is in demographic group $g$, and $k(\mu)$ was the average degree for the population over infectious period γ. For the PC, we used the $p_g$ that are calculated from the SB for both populations, assuming that they are the same (see Section 2 in Supplemental Methods).

We generated an expectation matrix $E$ to represent what we might expect mixing to look like over an infectious period given it occurs at random in the population. Each element of $E$ was calculated as:

$$e_{gh} = \frac{k_g(\mu)p_g}{k(\mu)} \frac{k_h(\mu)p_h}{k(\mu)}$$

Results for the comparison of $E$ to $M$ can be found in fig. S11 and S12.

**Section 5: Detailed methods for validating model results with 2013 DMV mortality data**

To show support for our models by comparing them to observed disease data, we obtained illness and mortality data available for bottlenose dolphins that washed ashore or were found floating, considered "stranded" from 2010-2015 from the National Marine Fisheries Service. We considered cases that occurred only in waters affected by the 2013 DMV epidemic (New Jersey, Delaware, Maryland, Virginia, North and South Carolina, Georgia and the counties of Nassau, Duval, St. John's, Flagler, Volusia and Brevard in Florida). We only considered mortalities that were not skeletal/mummified remains as the time of death for these can not be assumed to be during the epidemic. We also dropped all individuals for which age class was unknown or unrecorded. From the remaining data, we examined differences in mortality among age class (hereafter referred to as the "age data") and second, between adults of known sex, (hereafter referred to as the "sex data").

The DMV epidemic spanned 21 months (1-Jul-13 to 1-Mar-15), but not all recorded cases during this time period would be due to DMV [16]. Therefore, in order to obtain a better idea of the epidemic specific sickness and mortality, we removed the 'excess mortality' for each demographic group using methods adapted from [15]. For both the age and sex data we applied a negative binomial generalized linear model to the data with the number of cases as the response variable. We included a categorical predictor variable for the interaction between time period (pre DMV epizootic or during DMV epizootic) and age class (calf, juvenile, adult) or adult sex class (male, female), and state (fig. S13).

We observed that the number of recorded cases of calves before and during the epizootic were not significantly different (fig. S13). We also saw that in the state of Virginia the number of cases is lower compared to other states (fig. S13), but this effect was driven by the disproportionate number of unidentified age and sex cases compared to the other states. Therefore we did not consider cases from Virginia, or calves, for the remainder of the analysis.

From our glms, we found that adult dolphins stranded at an average rate of 3.1 and 18.1 cases per month, and juveniles at a rate of 1.7 and 12.1 cases per month before and during the epizootic respectively. We also found that adult males stranded at an average rate of 0.9 and 9.4 cases per month and adult females at a rate of 1.5 and 5.5 cases per month before and during the epizootic period respectively. Morris et al. assumed that the probability of a case recorded during the DMV epizootic time period being an actual DMV case is equal to the pre-epizootic case rate divided by the epizootic case rate for each age or sex class. Therefore, to remove excess mortality, we randomly removed a case from the data if it occurred between Jul-2013 and Mar-15, with this probability. To allow for variation in the final epidemic data, we repeated this process 1000 times to generate a confidence interval of epidemic mortality per age or sex class. We assumed that potential biases in the mortality data arising from non-uniform spatial distributions (e.g. from migration) or age and sex specific natural mortality risks are accounted for by removing these from the data.

To compare the final mortality case numbers for the age and sex class data to our simulation results we normalized the number of mortalities for each age or sex class to be an estimated proportion of recovered epidemic mortality for that age or sex class in the Atlantic dolphin populations. We assumed the same demographic group distribution used to generate our networks (see supplemental Section 2), and assumed the same population size used by Morris et al. of 26,317 individuals based on National Marine Fisheries Service stock assessment reports [46]. Because we found no information on how DMV mortality risk might vary based on age or sex, we also assumed that mortalities are proportional to the number of infected individuals in the population.

Finally, because the SB demographic distribution might not be representative of Atlantic bottlenose dolphins, and this distribution might also vary among the different Atlantic bottlenose dolphin stocks, we performed a sensitivity analysis where we calculated the proportion of each demographic group that stranded during the epizootic for various possible demographic distributions. For the age class comparison, we examined the proportion of juveniles to adults that stranded based on the following potential age class breakdowns:
10% of the population are juveniles, 90% are adults and calves;
20% of the population are juveniles, 80% are adults and calves;
30% of the population are juveniles, 70% are adults and calves;
40% of the population are juveniles, 60% are adults and calves.

For the adult sex class comparison, we considered the following potential sex class breakdowns:
35% of adults are male; 65% are female;
40% of adults are male; 60% are female;
45% of adults are male; 55% are female;
50% of adults are male; 50% are female;
55% of adults are male; 45% are female;
60% of adults are male; 40% are female;
65% of adults are male; 35% are female.

Results

For age class, we see that juveniles are still disproportionately found stranded compared to adults during the DMV outbreak for all hypothetical age class breakdowns (fig. S14; left). For adult sex class, we see that males are also still disproportionately found stranded compared to females during the DMV outbreak for all hypothetical sex class breakdowns (fig. S14; right). These results suggest that even if the demographic distribution of the Atlantic dolphin populations are different than that of SB, our model results are still supported by stranding data

**Section 6: Additional Disease Simulation Results**

Additional results can be found in fig. S15, S16 and S17.

**Section 7: PC Sensitivity Analysis**

Of the 99 focal individuals from the PC, 17 had a low confidence demographic group assignment. Of the 167 focal contacts from the PC, 84 had a low confidence demographic group assignment (See table S1). Therefore to account for potential errors in assigning demographics, we performed a sensitivity analysis in which the 5 low confidence adult males and 1 low confidence adult female are fully dropped from the analysis. Then, we randomly reassigned 33%, and then 66%, of the low confidence juveniles to either the adult male or adult female demographic group. Doing so would account for whether they were a misidentified young adult or older calf, as calves are included in the adult female group for our analysis. Then, we randomly reassigned 33%, and then 66%, of the low confidence contacts to another demographic group. We did not do a 100% reassignment of low confidence focals and contacts (a case representing us being wrong about every low confidence demographic assignment) because it left our sample size of juveniles too small to truly estimate differences.

We then re estimated our $k_g(D)$ and $M$ and regenerated 10 networks based on the new demographic degree distributions and demographic assortative mixing values. We reran our disease simulations on these networks to see if our epidemic outcomes were robust to potential errors in demographic assignment. We find that our epidemic outcomes held in both the 33% and 66% reassignment scenarios (fig. S18).

**Section 8: Estimating the effect of demographic specific degree and assortative mixing on infection results**

To examine whether demographic specific degree distributions or demographic assortativity is more important for understanding demographic specific risk, we also simulated a set of networks that had either null degree or null assortativity. To generate the null degree networks, we used demographic degree distributions that were calculated based on the population mean and variance (instead of the empirically estimated demographic specific values), but the demographic assortativity remained the same. To generate the null assortativity networks, we generated networks using empirical demographic-specific degree distributions, but assumed that assortativity is equal to our expectation matrix $E$. We then simulated disease spread on both sets of null networks and calculated the epidemic outcomes to compare to the empirical network simulations for both PC and SB. We compared these outcomes using a one-way ANOVA and Tukey's test.

**Section 9: Testing for differences in finer scale demographic groups using the SB**

We tested to see if there are differences in juvenile males and juvenile females, as well as between adult females and mother calf pairs using data from the SB, to see if our general demographic group assignments are justifiable.

For juveniles, we assessed the average synchrony degree for males and females over the average focal follow length for all follows on a juvenile individual. We used a generalized linear mixed model, where our response variable was the synchrony degree for each follow, with a categorical fixed effect for sex, and a continuous fixed effect for the length of the follow. We also included random effects for the year of the follow, and the ID of the dolphin to control for variability in data collected across years, and individual behavior of the dolphin respectively. We found that there is no difference in synchrony degree over the course of a focal follow for juvenile males and juvenile females (Supplementary fig. 21; left).

For adult females, we assessed how average synchrony degree varies based on whether or not they have a dependent calf present over the average focal follow length for all follows on an adult female individual. We used a generalized linear mixed model, where our response variable was the synchrony degree for each follow, with a categorical fixed effect for calf presence, and a continuous fixed effect for the length of the follow. We also included random effects for the year of the follow, and the ID of the dolphin to control for variability in data collected across years, and individual behavior of the dolphin respectively. We found that there is no significant difference in synchrony degree over the course of a focal follow for adult females when calves are present versus absent (Supplementary fig. 21; right).

### Appendix 1: Age-Sex Class and Mother-Calf Pair Assignment Rules for the Potomac-Chesapeake Dolphin Project

Developed by Ann-Marie Jacoby and Janet Mann

Hierarchy of Certainty (from highest to lowest): 1) Confirmed, 2) Probable, 3) Suspected
We consider Confirmed and Probable age and sex class assignments as “high confidence” for our analysis and Suspected age and sex class assignments as “low confidence”. If an individual has a high confidence adult age assignment but low confidence sex assignment then they are considered low confidence in their demographic group. See Supplemental Section 1 for how we divide individuals into demographic groups and handle “Unknown” age and sex class assignments.

**Important Definitions:**

**Infant Position:** “Infant swims under mother, melon or head lightly touches mother’s abdomen. Upon surfacing to breathe, the infant breaks infant position [for the duration of the surfacing], then angles back under the mother. Infant position swimming can be determined by the infant’s surfacing position slightly behind and </= 0.5m from the mother” [84].

**Echelon Position:** “Also referred to as contact swimming when the calf is close </= 30cm alongside the mother – roughly parallel, but sometimes slightly ahead, touching the mother’s flank intermittently or continuously above midline” (*84*).

**Sex Class Assignment:**

**Female, Confirmed (confirmed mother under criteria 3):**

Either:

1) Observation of genitals: short genital slit with two smaller mammary slits on either side and a small gap between the genital slit and anus [60,85,86].
2) Sexing data from biopsy sampling
3) Observed with a calf swimming in infant position or echelon [84] by a trained observer with one or more of the following conditions:
   Photograph Only:
   a) Photographed with a calf swimming in infant position or echelon for **three** independent surfacing bouts in **one survey or focal follow**. Photographs of the calf in infant position or echelon must be 5 minutes apart to count as an 'independent surfacing bout'. We use a value of 5 minutes to differentiate independent surfacing bouts of the calf in infant or echelon position, because the average infant position bout in Shark Bay is between 3.5-5.5 minutes depending on calf age.

b) Photographed with a calf swimming in infant position or echelon for **one** independent surfacing bout in each of **<u>three</u> observations (surveys or focal follow)**. Photographs of the calf in infant position or echelon must be 5 minutes apart to count as an 'independent surfacing bout'.
c) Photographed with a calf swimming in infant position or echelon for **one** surfacing bout that lasts for more than 5 minutes.

Live Observation Only:

d) A focal animal is observed with a calf swimming in infant position or echelon for **three** independent infant position or echelon bouts in **<u>one</u> focal follow**.
e) A focal animal is observed with a calf swimming in infant position or echelon for **one** independent infant position or echelon bouts in **<u>three</u> focal follows**.
f) A focal animal is observed with a calf swimming for **one** bout of infant position or echelon for greater than 5 minutes in **<u>one</u> focal follow**.
g) A non-focal animal or survey animal is observed with a calf swimming in infant position or echelon for **three** independent surfacing bouts in **<u>one</u> observation (either focal follow or survey, respectively)**.
h) A non-focal animal or survey animal is observed with a calf swimming in infant position or echelon for **one** independent surfacing bout in **<u>three</u> observations (either focal follow or survey, respectively)**.

Photograph & Live Observation:

i) An animal is photographed or observed with a calf swimming in infant position or echelon for **three** independent surfacing bouts or infant position/echelon bouts (focal follow only) within or **<u>across</u> observations (either survey or focal follow)**.

**Female, Probable (probable mother):**

Observed with a calf swimming in infant position or echelon [84] by a trained observer with one or more of the following conditions:

Photograph Only:

a) Photographed with a calf swimming in infant position or echelon for **two** independent surfacing bouts in **<u>one</u> observation (either survey or focal follow)**. Photographs of the calf in infant position or echelon must be 5 minutes apart to count as an 'independent surfacing bout'. We use a value of 5 minutes to differentiate independent surfacing bouts of the calf in infant or echelon position, because the average infant position bout in Shark Bay is between 3.5-5.5 minutes depending on calf age.
b) Photographed with a calf swimming in infant position or echelon for **one** independent surfacing bout in each of **<u>two</u> observations (either survey or focal follow)**.

Live Observation Only:

c) A focal animal is observed with a calf swimming in infant position or echelon for **one** infant position or echelon bout in each of **<u>two</u> focal follows**.

d) A focal animal is observed with a calf swimming in infant position or echelon for **two** independent infant position or echelon bouts in **__one__ focal follow**
e) A non-focal animal or survey animal is observed with a calf swimming in infant position or echelon for **one** independent surfacing bout in each of **__two__ observations (either focal follow or survey, respectively)**.
f) A non-focal animal or survey animal is observed with a calf swimming in infant position or echelon for **two** independent surfacing bouts in **__one__ observation (either focal follow or survey, respectively)**.

Photograph & Live Observation:

j) An animal is photographed or observed with a calf swimming in infant position or echelon for **two** independent surfacing bouts or infant position/echelon bouts (focal follow only) within or **__across__ observations (either survey or focal follow)**.

**Female, Suspected (suspected mother):**

Observed with a calf swimming in infant position or echelon [84] by a trained observer with one or more of the following conditions:

Photograph Only:

a) Photographed with a calf swimming in infant position or echelon for **one** independent surfacing bout in **__one__ survey or focal follow**. Photographs of the calf in infant position or echelon must be 5 minutes apart to count as an 'independent surfacing bout'. We use a value of 5 minutes to differentiate independent surfacings bouts of the calf in infant or echelon position (average infant position bout in SB is between 3.5-5.5 minutes depending on calf age).

Live Observation Only:

b) A focal animal observed with a calf swimming in infant position or echelon for **one** infant position or echelon bout during **__one__ focal follow**.
c) A non-focal animal or survey animal is observed with a calf swimming in infant position or echelon for **one** independent surfacing bout in **__one__ observation (either focal follow or survey, respectively)**.

Photograph & Live Observation:

d) An animal is photographed or observed with a calf swimming in infant position or echelon for **one** independent surfacing bout or infant position/echelon bout (focal follow only) within or **__across__ observations (either survey or focal follow)**.

**Male, Confirmed- High Confidence:**

Either:

1) Observation of genitals: (a) erection; (b) long genital slit and large gap between the genital slit and anus [60,86]; (c) no mammary slits
2) Sexing data from biopsy sampling

**Male, Probable- High Confidence:**
Must have *three* of the criteria described in the male suspected category (below).

**Male, Suspected- Low Confidence:**
Either:
1) Observed body size of a full grown adult male (average Sarasota, Florida: 265 cm and 150cm wide [87]); determined during survey and focal follow
   a) Sexual dimorphism is apparent in PC animals, with the greatest male length off of North Carolina being 334 cm and female length being 285 cm [86].
2) Not observed with a dependent calf throughout the duration of the sampling period.
3) Behavior: Exhibits multi-year associations with a probable male (see above). For animals to be associated with one another they have to be documented in the same survey or focal follow. Therefore multi-year associations mean animals were seen in the same survey(s) or focal follow(s) in one or more years.
4) Behavior: Mounting or sex play by a confirmed, probable, or suspected adult animal based on size
   a) Mounting by adult females is exceedingly rare [88]
5) Synchronous displays by confirmed, suspected, or probable adult animals [25,60].
   a) Synchronous displays are defined as: Surface displays that involve slapping a body part (belly, face, jaw, pectoral fin, dorsal side) on the water surface; spy hops (bring head vertically out of water perpendicular to water surface; and porp outs at the same time as and in close proximity to another individual.
   b) This does <u>not</u> include breathing synchrony as this would bias our analysis

**Unknown Sex:**
Dolphins that cannot be assigned as either a suspected, probable or confirmed female or male due to a lack of data are assigned an unknown sex.

**<u>Age Class Assignment:</u>**
**Adult, Confirmed- High Confidence:**
A female that has a dependent calf (see female and calf assignment criteria as well as mother-calf pair rules).

**Adult, Probable- High Confidence:**
More than one sighting of an animal described as a suspected adult using criteria 1 and 2.

**Adult, Suspected- Low Confidence:**
Either:

1) Observed body size of a full grown adult male (average Sarasota, Florida: 265cm and 150cm wide) or female (average Sarasota, Florida: 249cm long and 140cm wide [87]; determined during a surveys and focal follows by a trained observer
2) Determined in photographs by looking for body and fin size relative to other animals in the survey of known or noted age (determined post-hoc by a trained photo-ID coder)
3) Is either an adult or a juvenile based on: a) observations made by a trained observer during a survey or focal follow, or b) the animal's body and fin size in photographs relative to other animals in the survey or focal follow (determined post-hoc by a trained photo-ID coder). The animal is assigned both Suspected Adult and Suspected Juvenile, which is also categorized with the code "T2"

**Juvenile, Confirmed- High Confidence:**

Previously observed as a calf (see calf aging rules) and is weaned An animal is considered weaned if it: a)is not observed in infant position during data collection or from photographs, b) sightings with its mother decline to <50% throughout a sampling year, and/or c) its mother is observed during data collection or from photographs with a new calf that is younger in age.

**Juvenile, Probable- High Confidence:**

More than one observation of an animal in a survey or focal follow described as a suspected juvenile using criteria 1) and 2)

**Juvenile, Suspected- Low Confidence:**

Either:

1) Observed body size: ¾ or less the size of an adult and is not observed in infant position; determined during survey and focal follow by a trained observer
    a) The minimum weaning date of coastal bottlenose dolphins in Sarasota Bay, FL is between 1 and 2 years of age [89]. Using the growth curve [87], animals this age are a length of ~185cm, which is ¾ the size of an adult female.
2) Determined in photographs looking for body and fin size relative to others in the survey of known or noted age (determined post-hoc by a trained photo-ID coder)
3) Is either a juvenile or an adult based on: a) observations made by a trained observer during a survey or focal follow, or b) the animal's body and fin size in photographs relative to other animals in the survey or focal follow (determined post-hoc by a trained photo-ID coder). The animal is assigned both Suspected Juvenile and Suspected Adult, which is also categorized with the code "T2"
4) Is either a juvenile or an older calf based on: a) observations made by a trained observer during a survey or focal follow, b) the animal's body and fin size in photographs relative to other animals in the survey or focal follow (determine post-hoc by a trained photo-ID coder), or c) having a "low" or "suspected" calf assignment. The animal is assigned both Suspected Calf and Suspected Juvenile, which is also categorized with the code "T1"

**Calf, Confirmed:**

One or both:

1) Exhibits behaviors and/or physical features of a newborn (0-3 months or 0-91.5 days), which include the following:
   a) Features at 1 month (<30.5 days):
      - 95 and 134cm (at birth);
      - flopped over dorsal fins (<1 day) [90,91];
      - whiskers (<4 days) [84,92];
      - fetal folds and lumpiness or fetal lines and visible neck (<7 days) [84];
      - cork-up surfacings (<7 days) [84];
      - 30-100% chin-up surfacings (<14 days) [84];
      - 2:1 echelon to infant position ratio [84]
   b) Features at 2-3 months (30.5-91.5 days):
      - <10% chin-up surfacings (30.5-61 days) [84];
      - 1:2 echelon to infant position ratio (30.5-61 days) or infant position only [84];
      - faint fetal lines (30.5-91.5 days) [84]
2) A newborn (criteria 1) or a calf this is not a newborn (>3 months, has no fetal lines, <185cm, see Body Size Scale for details) observed in infant position or echelon (newborn only) with an animal (i.e, the presumed mother - the same animal within and across observations) by a trained observer with one or more of the following conditions:
   Photograph:
   a) Photographed swimming in infant position or echelon with the presumed mother for **three** independent surfacing bouts in **<u>one</u> survey or focal follow**. Photographs of the calf in infant position or echelon must be 5 minutes apart to count as an 'independent surfacing bout'. We use a value of 5 minutes to differentiate independent surfacing bouts of the calf in infant or echelon position, because the average infant position bout in Shark Bay is between 3.5-5.5 minutes depending on calf age.
   b) Photographed swimming in infant position or echelon with the presumed mother for **one** independent surfacing bout in each of **<u>three</u> observations (surveys or focal follow)**. Photographs of the calf in infant position or echelon must be 5 minutes apart to count as an 'independent surfacing bout'.
   c) Photographed swimming in infant position or echelon with the presumed mother for **one** surfacing bout that lasts for more than 5 minutes.

   Live Observation:
   d) A focal animal is observed swimming in infant position or echelon with the presumed mother for **three** independent infant position or echelon bouts in **<u>one</u> focal follow**.
   e) A focal animal is observed swimming in infant position or echelon with the presumed mother for **one** independent infant position or echelon bouts in **<u>three</u> focal follows**.

f) A focal animal is observed swimming in infant position or echelon with the presumed mother for **one** bout of infant position or echelon for greater than 5 minutes in **one focal follow**.
g) A non-focal animal or survey animal is observed swimming in infant position or echelon with the presumed mother for **three** independent surfacing bouts in **one observation (either focal follow or survey, respectively)**.
h) A non-focal animal or survey animal is observed swimming in infant position or echelon with the presumed mother for **one** independent surfacing bouts in **three observations (either focal follow or survey, respectively)**.

Photograph and Live Observation:

i) An animal is photographed or observed swimming in infant position or echelon with the presumed mother for **three** independent surfacing bouts or infant position/echelon bouts (focal follow only) within or **across observations (either survey or focal follow)**.

**Calf, Probable:**
A calf that is not a newborn (>3 months, has no fetal lines, >185cm, see Body Size Scale for details) observed in infant position (not echelon) with an animal (i.e, the presumed mother - the same animal within and across observations) by a trained observer with one or more of the following conditions:
Photograph:

a) Photographed swimming in infant position with the presumed mother for **two** independent surfacing bouts in **one observation (either survey or focal follow)**. Photographs of the calf in infant position must be 5 minutes apart to count as an 'independent surfacing bout'. We use a value of 5 minutes to differentiate independent surfacing bouts of the calf in infant or echelon position, because the average infant position bout in Shark Bay is between 3.5-5.5 minutes depending on calf age.
b) Photographed swimming in infant position with the presumed mother for **one** independent surfacing bout in each of **two observations (either survey or focal follow)**.

Live Observation:

c) A focal animal is observed swimming in infant position with the presumed mother for **two** independent infant position or echelon bouts in **one focal follow**.
d) A focal animal is observed swimming in infant position with the presumed mother for **one** independent infant position or echelon bouts in **two focal follows**.
e) A non-focal animal or survey animal is observed swimming in infant position or echelon with the presumed mother for **two** independent surfacing bouts in **one observation (either focal follow or survey, respectively)**.
f) A non-focal animal or survey animal is observed swimming in infant position or echelon with the presumed mother for **one** independent surfacing bouts in **two observations (either focal follow or survey, respectively)**.

Photograph & Live Observation:

g) An animal is photographed or observed swimming in infant position with the presumed mother for **two** independent surfacing bouts within or infant position bouts (focal follow only) **<u>across</u> observations (either survey or focal follow)**.

**Calf, Suspected:**
Either:

1) Does not exhibit physical or behavioral features of a newborn or older calf beyond its small size relative to others observed during data collection or in photographs and was noted as a calf by a trained observer or photo-ID coder
2) A calf that is not a newborn (>3 months, has no fetal lines, >185cm, see Body Size Scale for details) observed in infant position (not echelon) with an animal (i.e, the presumed mother) by a trained observer with one or more of the following conditions:
   Photograph:
   a) Photographed swimming in infant position with the presumed mother for **one** independent surfacing bout in **<u>one</u> survey or focal follow**. Photographs of the calf in infant position must be 5 minutes apart to count as an 'independent surfacing bout'. We use a value of 5 minutes to differentiate independent surfacings bouts of the calf in infant or echelon position (average infant position bout in SB is between 3.5-5.5 minutes depending on calf age).

   Live Observation:
   b) A focal animal is observed swimming in infant position with the presumed mother for **one** infant position or echelon bout in **<u>one</u> focal follow**.
   c) A non-focal animal or survey animal is observed swimming in infant position with the presumed mother for **one** independent surfacing bout in **<u>one</u> observation (either focal follow or survey, respectively).**

   Photograph & Live Observation:
   d) An animal is photographed or observed swimming in infant position with the presumed mother for **one** independent surfacing bout or infant position bout (focal follow only) within or **<u>across</u> observations (either survey or focal follow)**.

**Unknown Age**
Dolphins that cannot be assigned as a calf or as a suspected, probable or confirmed juvenile or adult due to a lack of data are assigned an unknown age.

**<u>Mother-Calf Pair Assignment:</u>**
A female and calf are determined using the behaviors and physical features listed in the female and calf assignment rules above.

**Mother-Calf Pair, Confirmed:**
If an animal that has been assigned as a confirmed calf and an animal that has been assigned as a confirmed mother (see above) are observed together with the calf in echelon or infant position with the mother, then they are a confirmed mother-calf pair.

**Mother-Calf Pair, Probable:**

If an animal has been assigned as a probable calf and an animal that has been assigned as a probable mother (see above) are observed together with the calf in echelon or infant position with the mother, then they are a probable mother-calf pair.

**Mother-Calf Pair, Suspected:**

If an animal that has been assigned as a suspected calf and an animal that has been assigned as a suspected mother (see above) are observed together with the calf in infant position with the mother, then they are a suspected mother-calf pair.

### Supplemental Figures

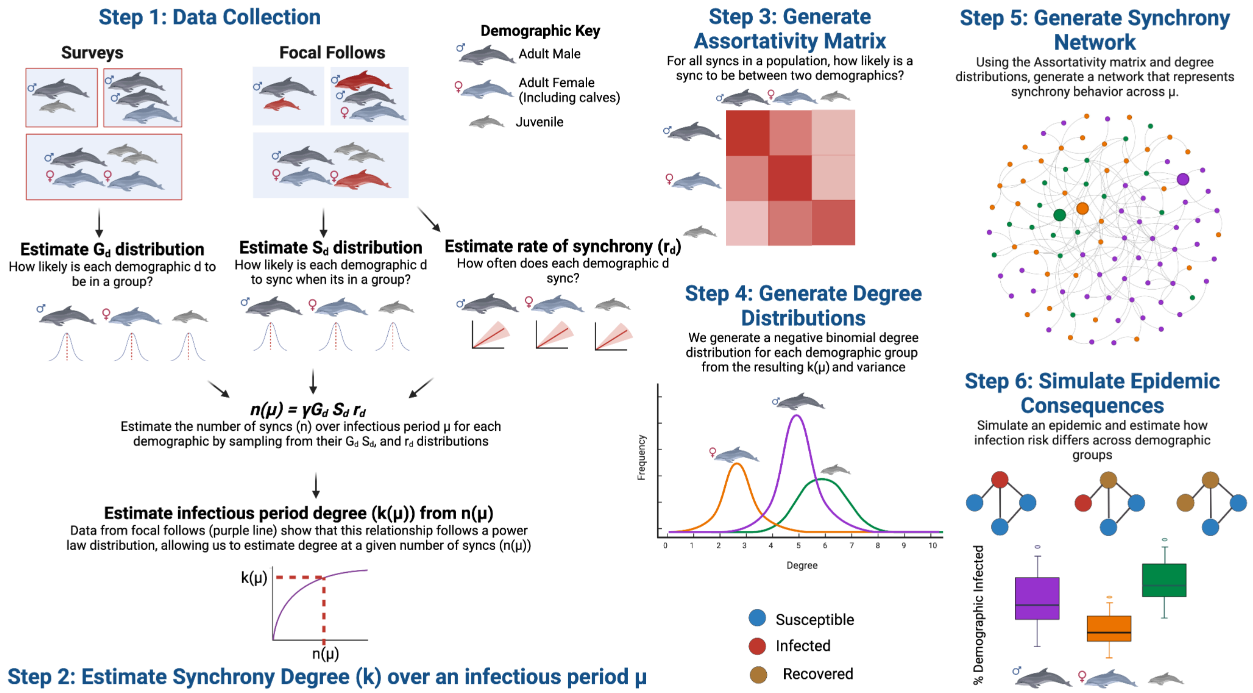


**Fig. S1:** Summary of full methods (expanded from main text Fig. 1). Created with BioRender.com

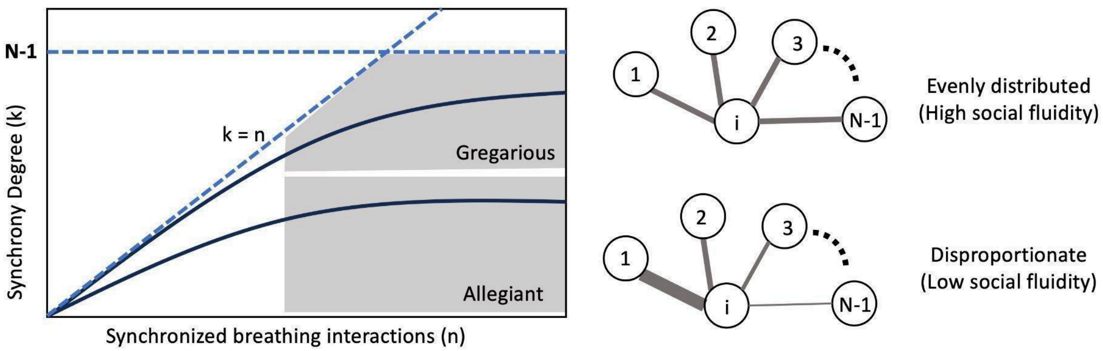


Fig. S2: Representation of the social fluidity of synchronized breathing contacts in bottlenose dolphins, adapted from Colman et al., 2021 Fig. 1. Dashed blue lines mark the boundary of the region where data points can feasibly be found for a population of size N. The mean degree is plotted representing either gregarious or allegiant social behavior. As the number of observed synchronized breathing interactions grows, synchrony degree increases; the rate at which it increases is known as social fluidity, and influences how we categorize synchronized breathing, and how we can estimate synchrony degree at a higher number of synchrony interactions. The weight (number) of the interactions between individual i and their contacts represents the propensity of i to interact with each of the other individuals in the group.

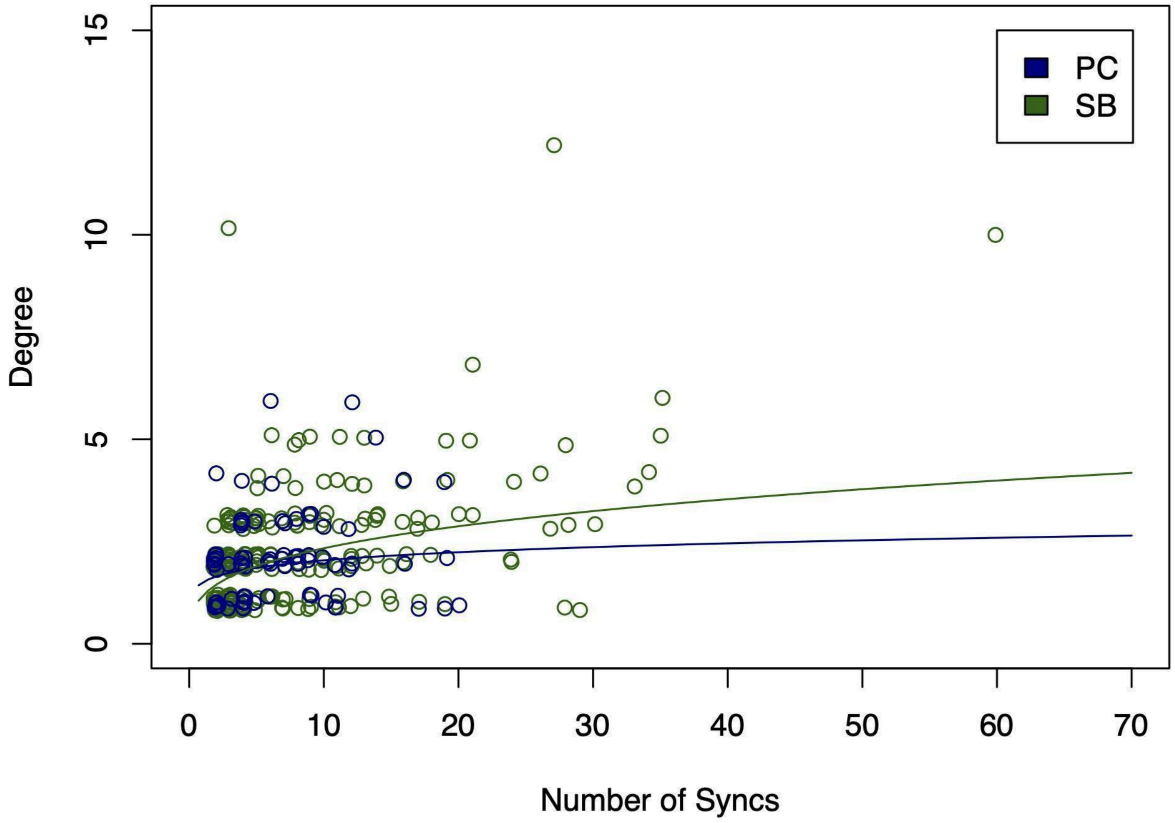


**Fig. S3:** The power law curves fitted to the PC (blue) and SB (green) focal follow data respectively. We found that most of the divergence occurs when the PC becomes sparse (high number of syncs) due to shorter follow lengths in the PC.

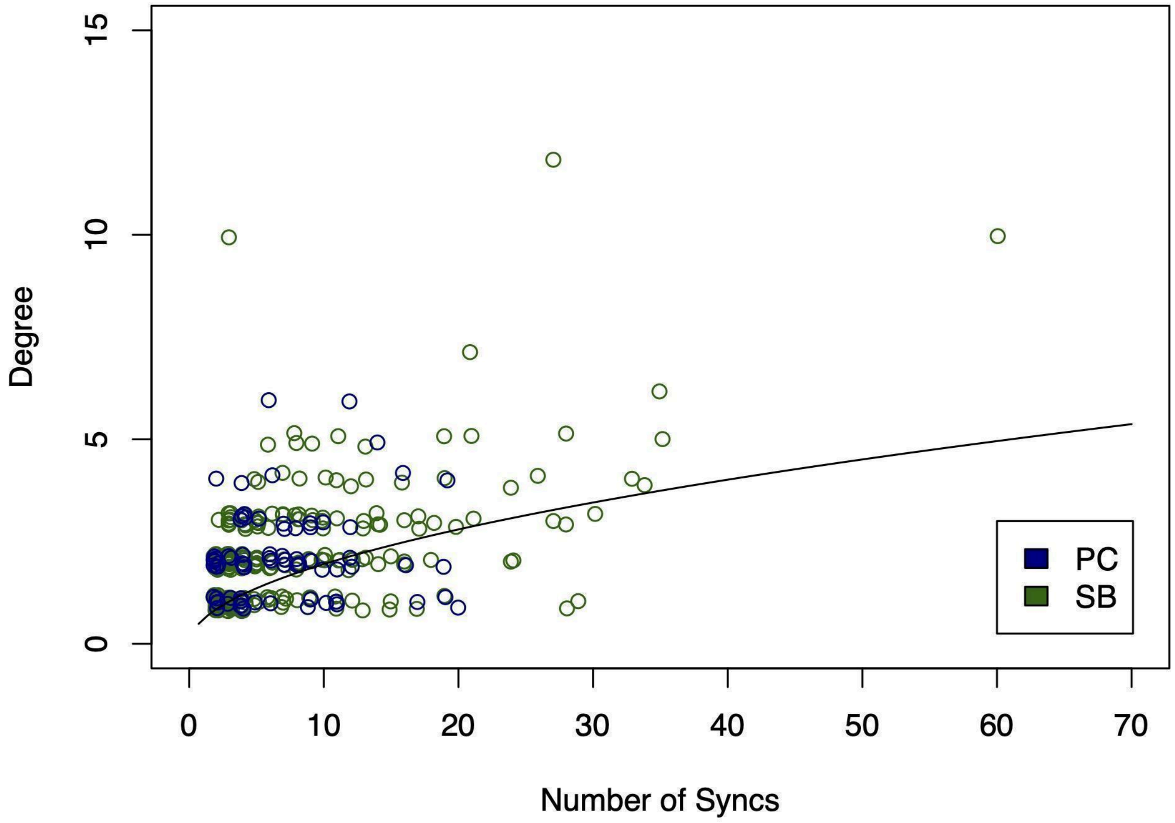


**Fig. S4:** Because of the similar curves generated using PC and SB data, we assumed that the relationship between degree and number of syncs was consistent across the two populations, and calculated one curve to establish this relationship and our α and λ.

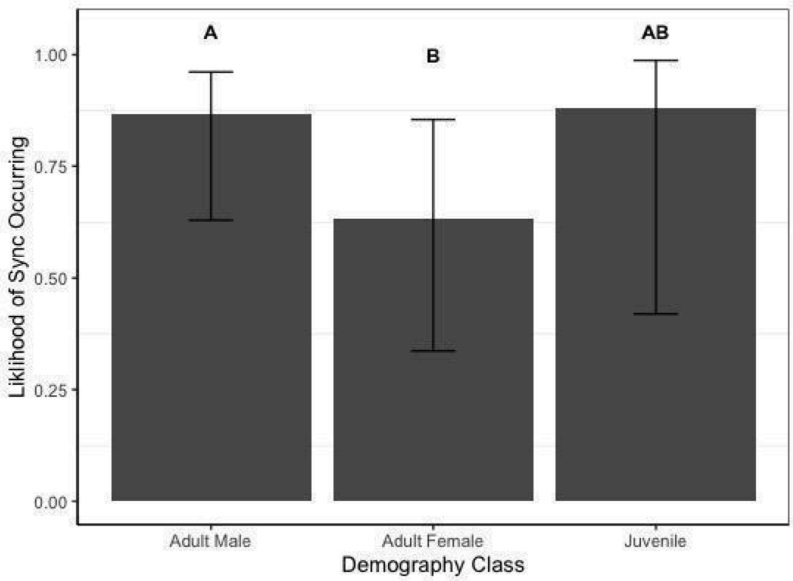


**Fig. S5:** The $S_d$ for the PC by demographic group. We find that males are more likely to sync than females, and that juveniles also have a high likelihood of syncing, but there is more variability in their $S_d$ values.

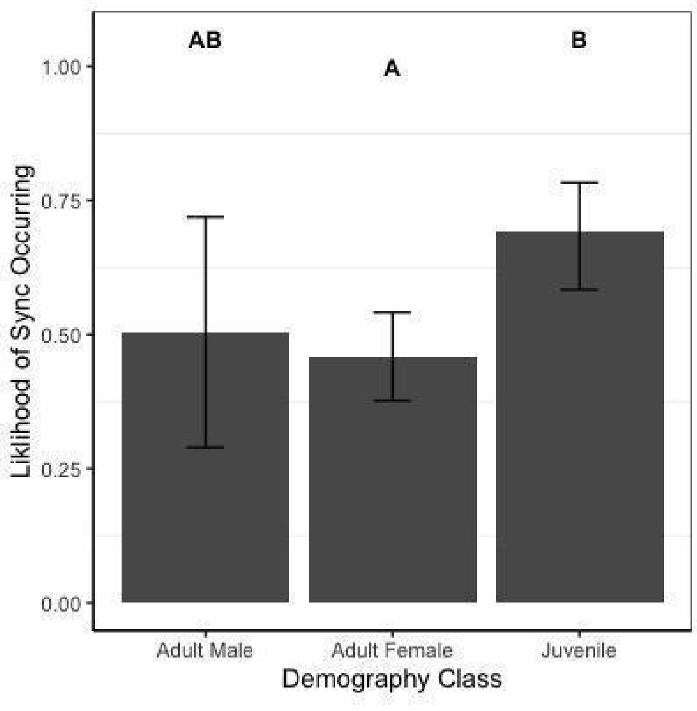


**Fig. S6:** The $S_d$ for the SB by demographic group, where letters indicate significance. We find that juveniles are more likely to sync than adult females, and that adult males also have a high likelihood of syncing, but there is more variability in their $S_d$ values.

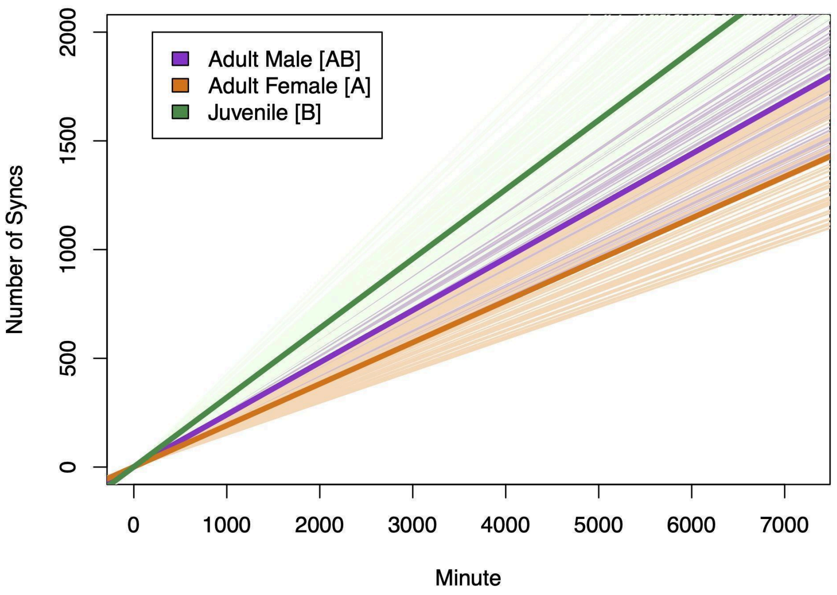


**Fig. S7:** The estimated $r_d$ for each demographic group in the PC. Letters in the legend indicate significance. We find that juveniles have the highest rate of synchrony and adult females have the lowest, and that adult males are more variable in their sync rates.

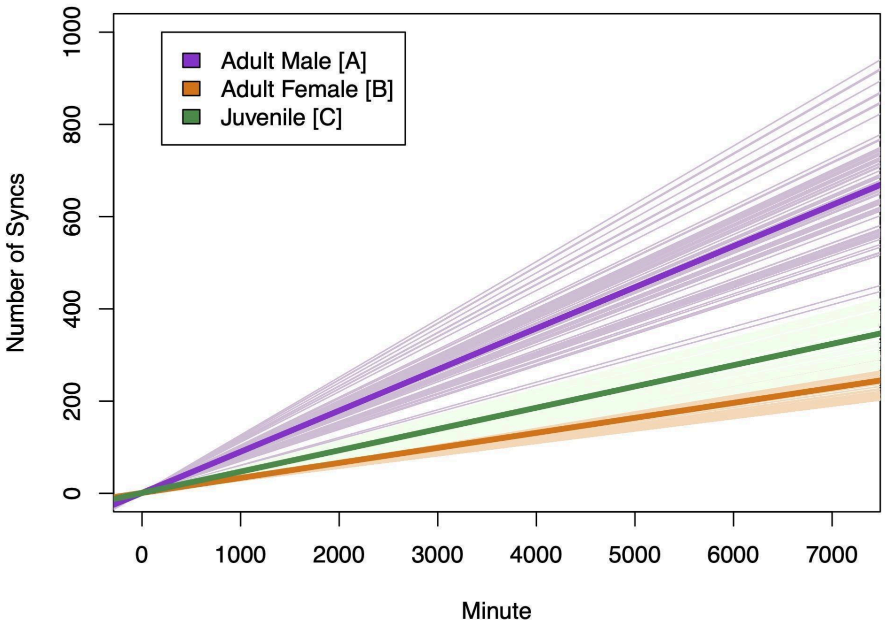


**Fig. S8:** The estimated $r_d$ for each demographic group in the SB. Letters in the legend indicate significance. We find that adult males have the highest rate of synchrony followed by juveniles, and adult females have the lowest.

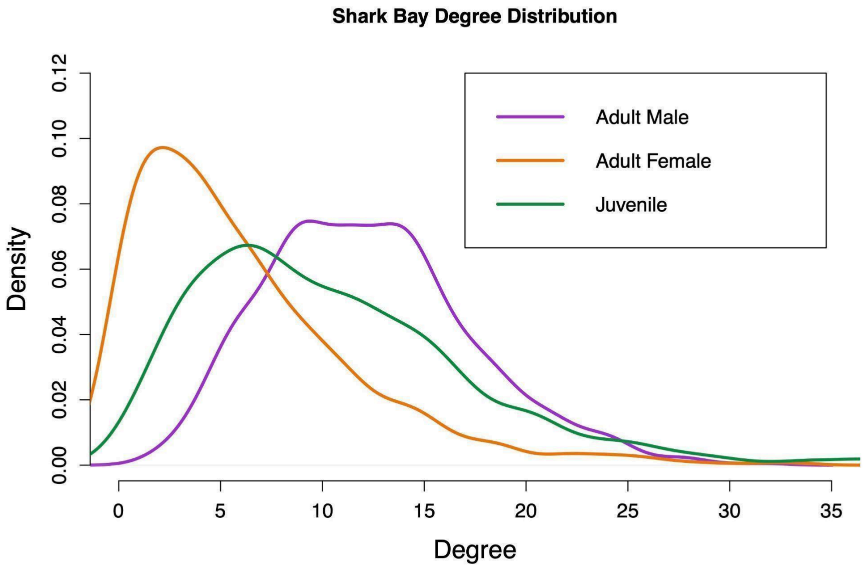


**Fig. S9:** The estimated degree distribution for the SB over a DMV infectious period. We find that females have the smallest average degree, followed by juveniles, and adult males have the largest average degree.

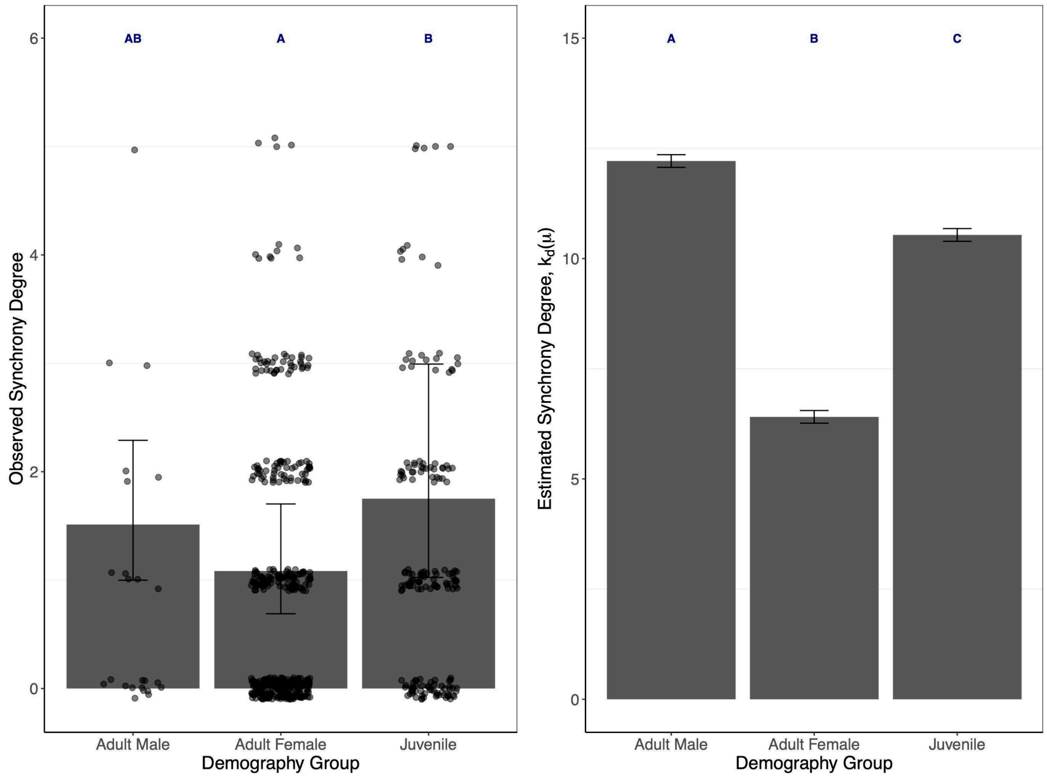


**Fig. S10:** The estimated degree over a DMV infectious period γ for each demographic group ($k_d(\mu)$) in the SB (left) compared to the estimated average degree over the average SB follow ($k_d(\mu)$) (right). Letters indicate significance. **We see that the overall demographic specific trends are maintained when we use our methodology to estimate degree over a DMV infectious period.**

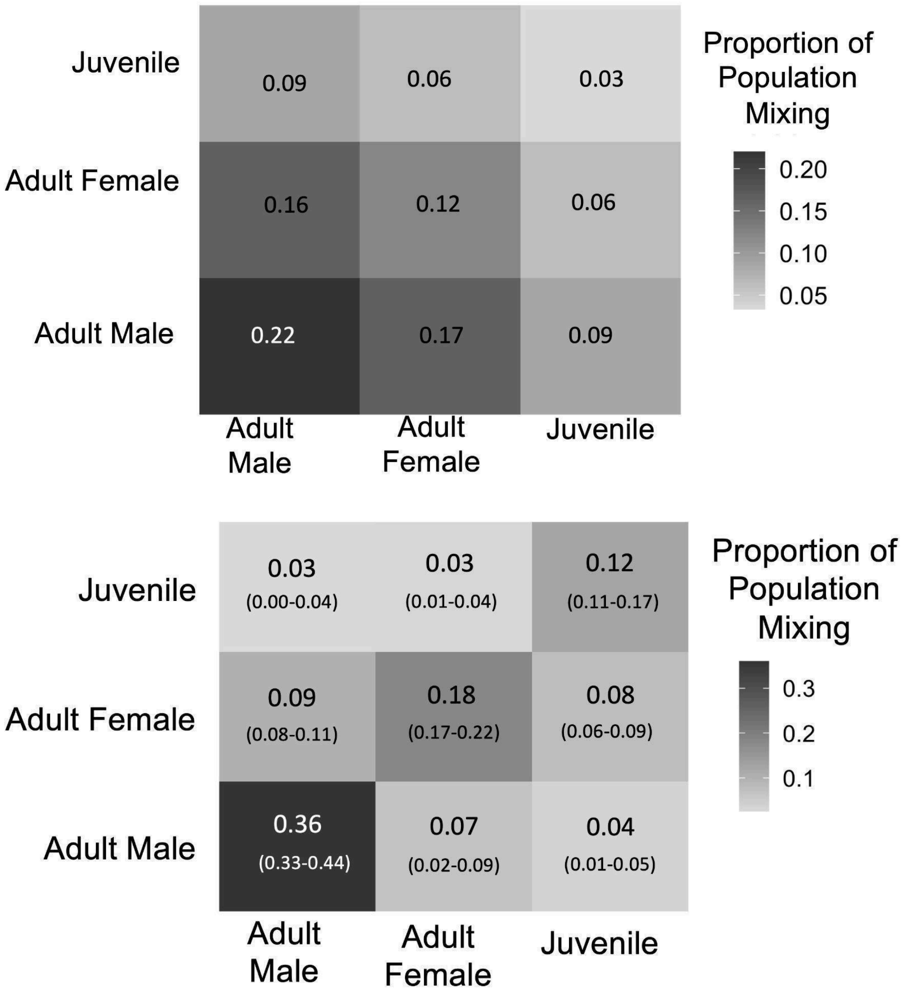


**Fig. S11:** The expectation mixing matrix E (top) compared to the PC empirical mixing matrix M (bottom). We see that our predicted values and confidence intervals of mixing are higher along the diagonal than the expected mixing values if mixing was completely random. This suggests **that all demographics are more assortative than what would be expected at random indicating preferential contacts within their own demographic groups.**

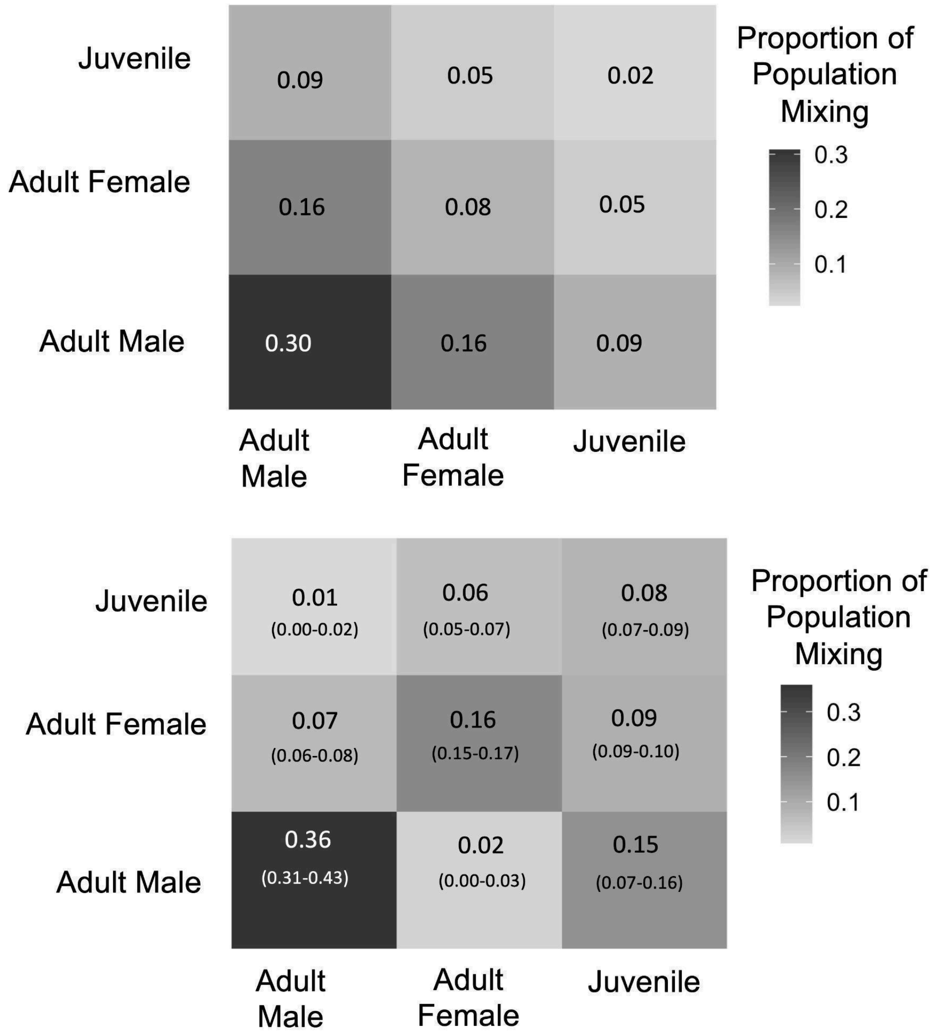


**Fig. S12:** The expectation mixing matrix E (top) compared to the SB empirical mixing matrix M (bottom). We see that our predicted values and confidence intervals of mixing are higher along the diagonal than the expected mixing values if mixing was completely random. This suggests **that all demographics are more assortative than what would be expected at random indicating preferential contacts within their own demographic groups.**

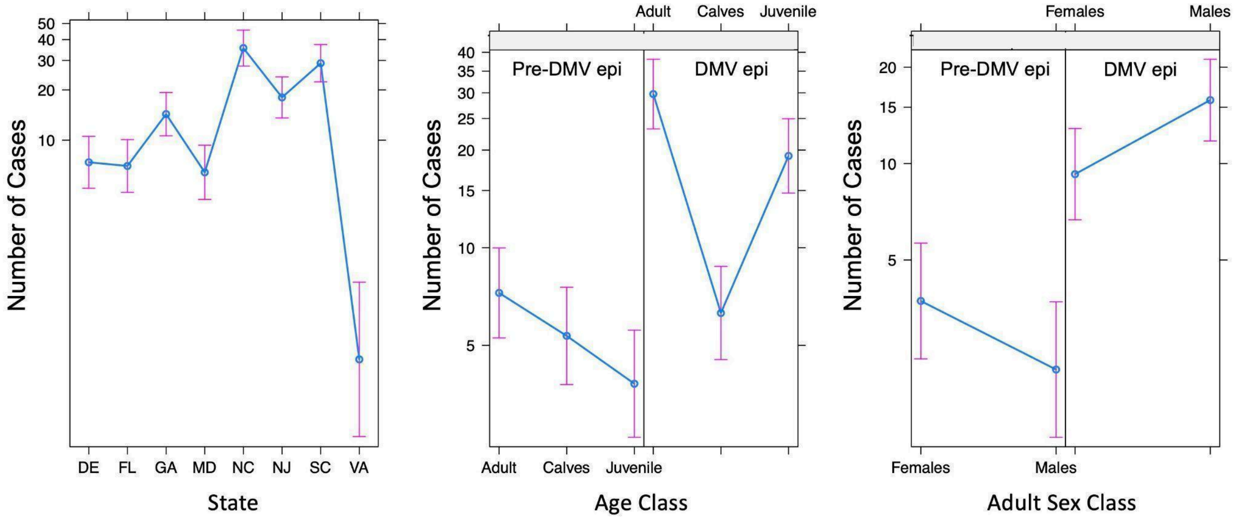


**Fig. S13:** The glm predictions for number of cases by state between 2010-2015 (left), and for age (middle) and adult sex (right) class before and during the DMV epizootic. We considered the average number of pre DMV cases by age and sex class as a measure of excess mortality, and removed this number of mortalities from the DMV epizootic data. We found no significant difference between calf recovery numbers in these time periods (i.e. no excess mortality). We also found that the number of cases is lower in Virginia compared to other states, but this was driven by the small sample size of individuals of known age class. Therefore we did not consider cases from the state of Virginia or calves in our comparison.

.

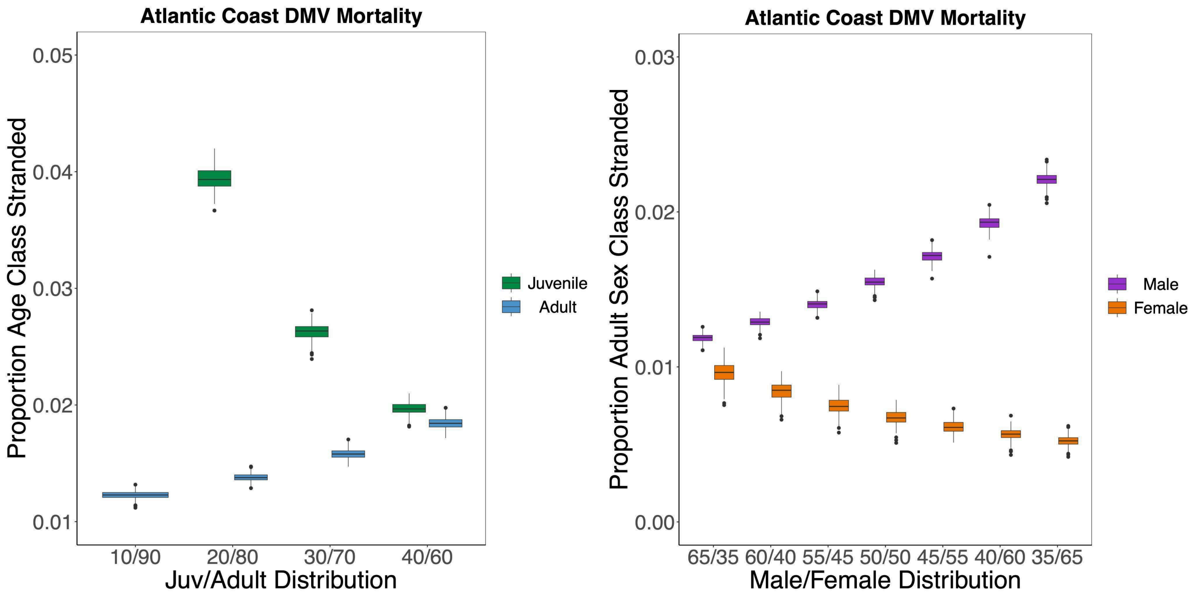


**Fig. S14:** The proportion of each age class (left) and adult sex class (right) that stranded during the 2013 DMV epidemic for different age and sex class distributions.

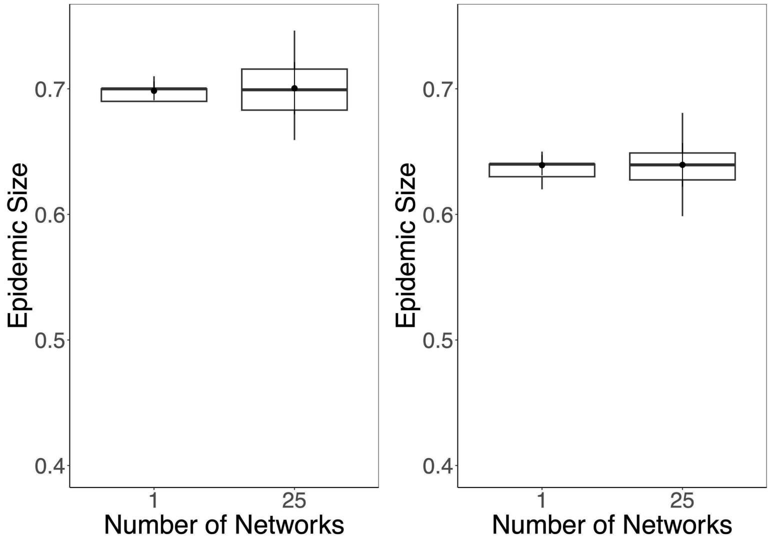


**Fig. S15**: Epidemic outcomes are not different in one network compared to across networks for both the PC (left) and SB (right), demonstrating that 25 generated networks is enough to capture contact variability.

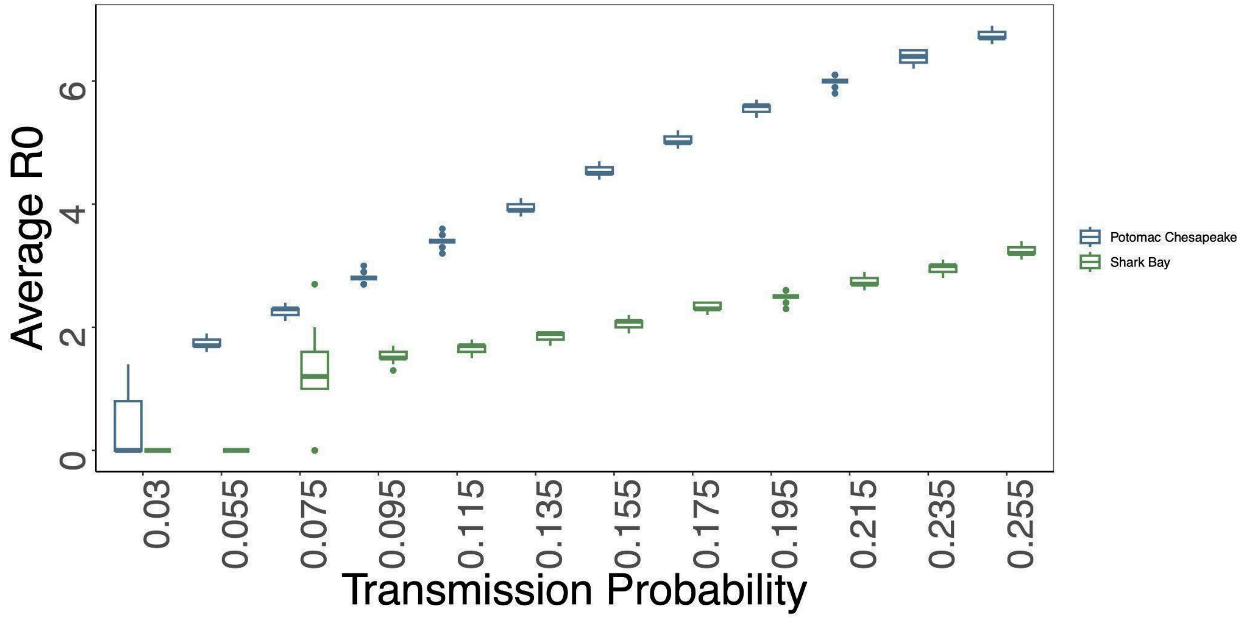


**Fig. S16:** The transmission probabilities required to achieve certain R0 values in our disease simulations for the PC (blue) and SB (green) networks. **We see that to achieve an R0 of 1.9, we need an transmission probability of 0.055 and 0.135 in the PC and SB respectively.**

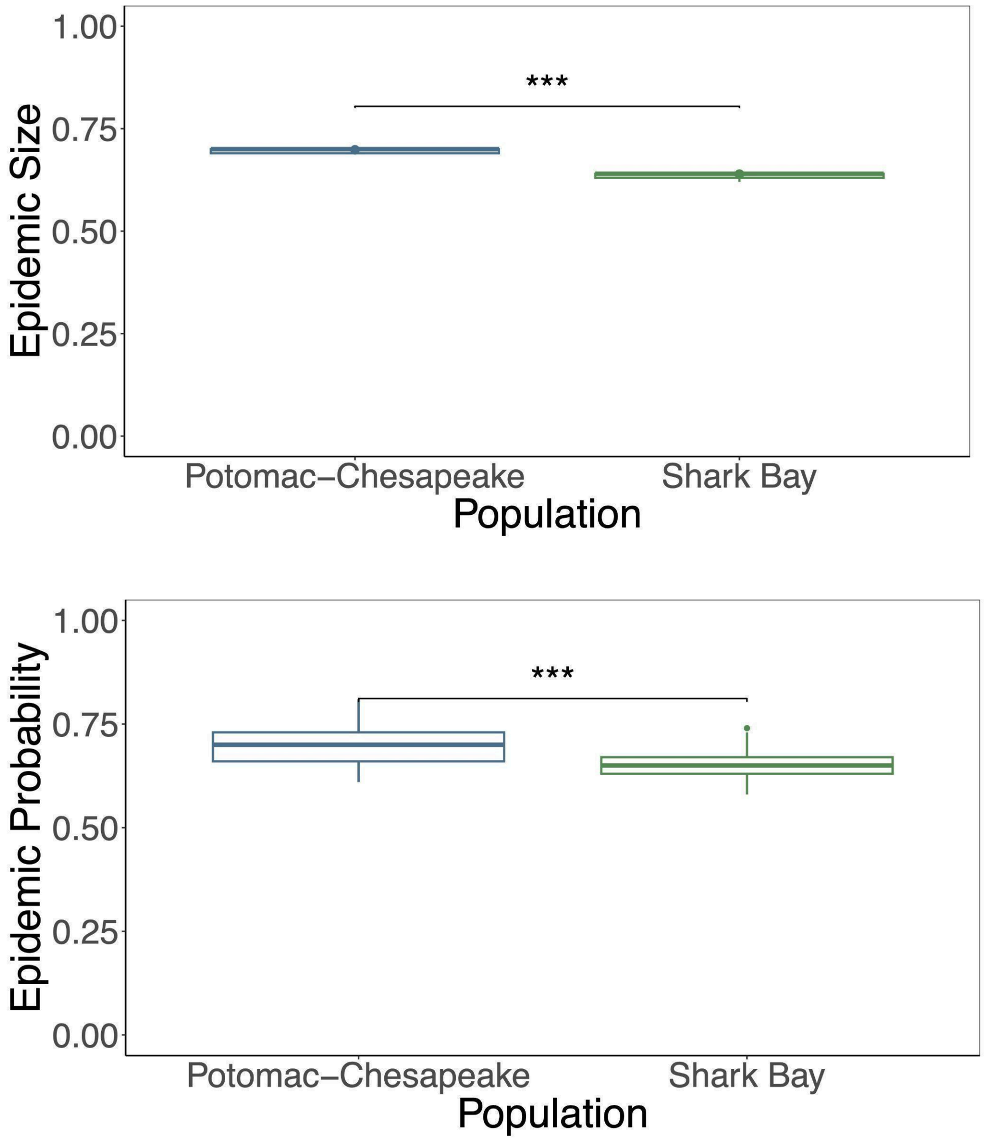


**Fig. S17:** The average epidemic size (top) and probabilities (bottom) for a simulated DMV epidemic with an R0 of 1.9 for the PC and SB. **Epidemic sizes and probabilities are higher in the PC networks compared to the SB networks**. At an R0 of 1.9, epidemic size (PC= 69.8%, SB = 63.9%, p = <0.0001) and probability (PC = 70.16%, SB= 64.9%, p=0.0002) is significantly higher in the PC compared to the SB

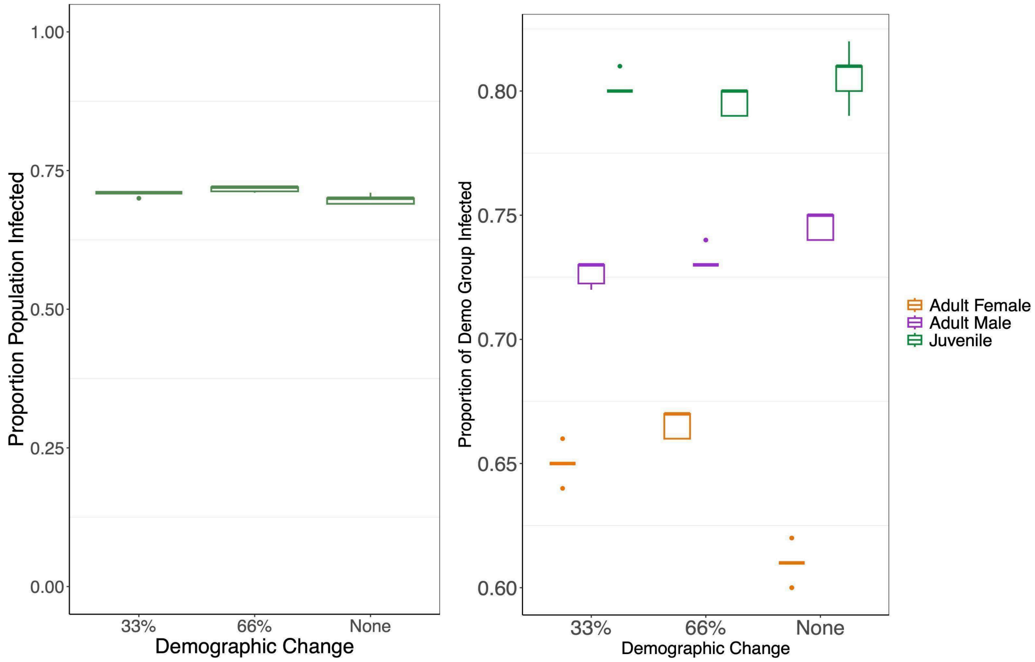


**Fig. S18:** The epidemic sizes (left) and proportion of each demographic group infected (right) when 33% and 66% of individuals with low confidence demographic group assignments were changed to another demographic group, compared to our empirical results (labeled "None"). We see that there are no changes to overall population impact for the PC dolphins, or demographic specific risks when 66% of low confidence individuals are assigned to a different demographic group, **indicating that our results are robust to potential the misidentification of individuals' age and sex classes.**

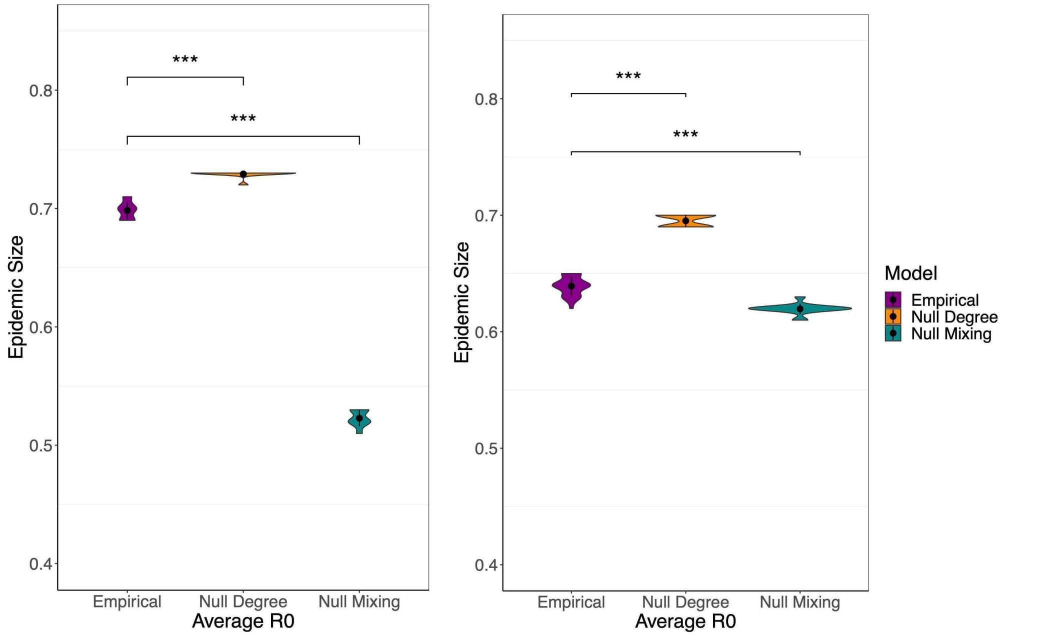


**Fig. S19:** The average epidemic sizes for PC (left) and SB (right) simulated DMV epidemics with an R0 of 1.9. We find that in the null degree scenario, epidemic size is higher than in our empirical model. In the null mixing scenario epidemic sizes were lower. **This indicates that demographic assortative mixing drives population level epidemic risks.**

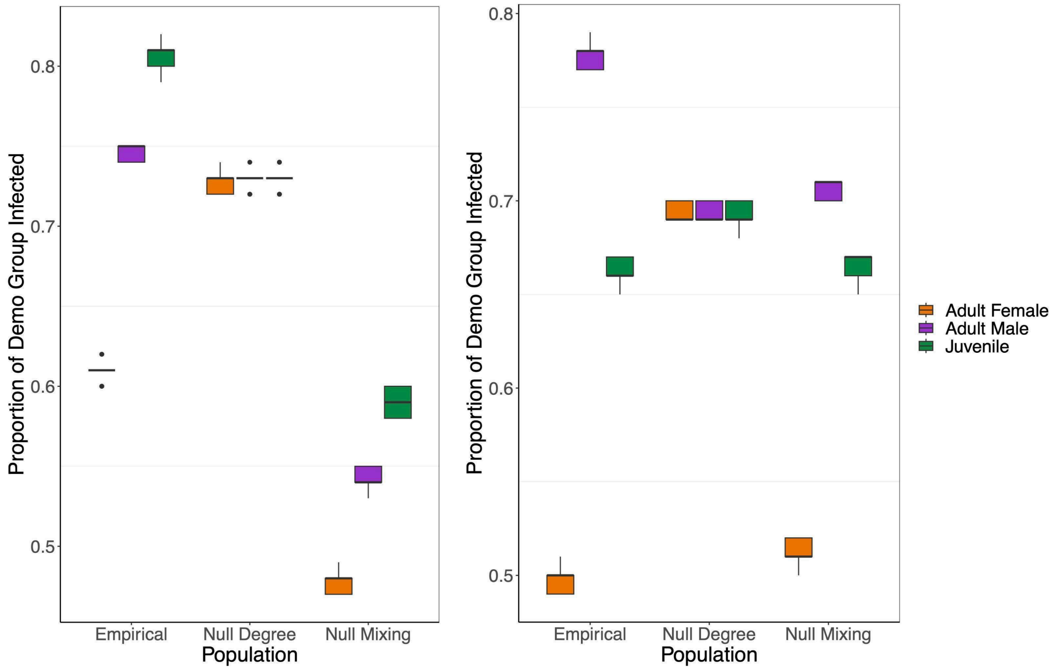


**Fig. S20:** The average proportion of each demographic group infected for PC (left) and SB (right) simulated DMV epidemics with an R0 of 1.9. We find that in the null degree scenario, we lose our demographic specific infection risks in both populations, but it is maintained in the null mixing scenarios. **This indicates that demographic specific degree drives demographic specific epidemic risks.**

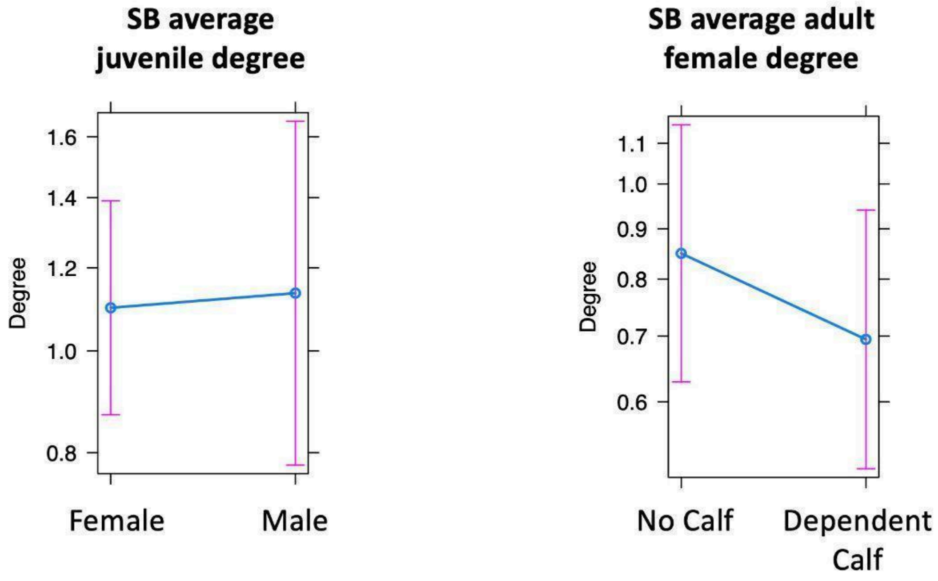


**Fig. S21:** Estimated synchrony degree over an average focal follow (~2 hours) using the SB data. We find that juvenile **degree does not vary significantly by sex**. **We also find that degree does not vary significantly when a calf is present versus absent for adult females**.

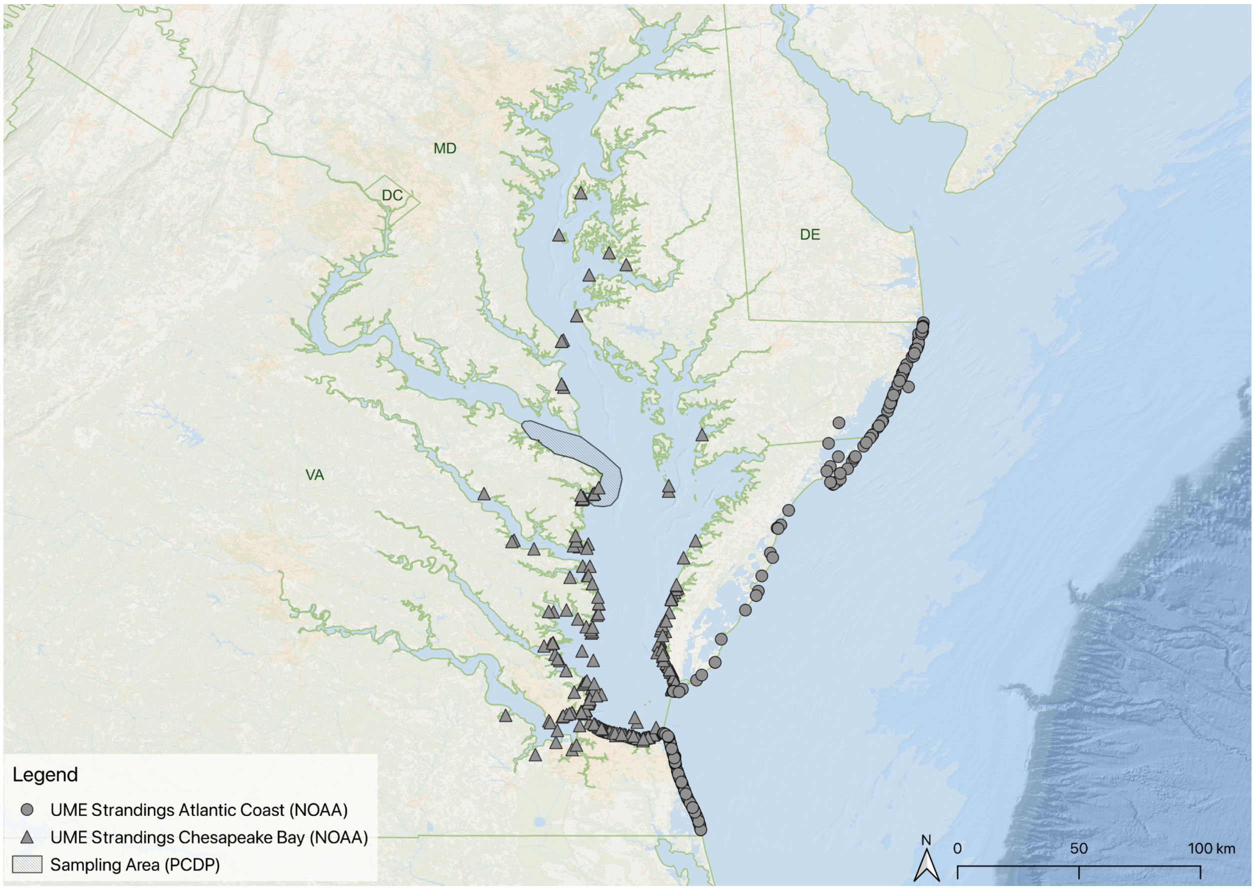


**Fig. S22.** The plurality of strandings during the 2013-2015 unusual mortality event (UME) associated with DMV occurred in Virginia and Maryland waters. The majority of these strandings occurred in the Chesapeake Bay (triangles). The PC study area is shaded in gray.

### Supplementary Tables

**Table S1:** Demographic group assignments and confidences of both focal individuals and their contacts for both the PC and the Shark Bay Dolphin Research Project (SB). Confidences for the PC were determined using Appendix 1, and methodology from supplementary Section 1 and main text Section 3.1.1.

| Population | Demographic Group | Total # of High Confidence Focals | Total # of Low Confidence Focals | Total # of High Confidence Contacts | Total # of Low Confidence Contacts |
|---|---|---|---|---|---|
| PC | Adult Male | 38 | 5 | 54 | 26 |
| PC | Adult Female | 40 | 1 | 23 | 22 |
| PC | Juvenile | 4 | 11 | 6 | 36 |
| SB | Adult Male | 25 | NA | 104 | 10 |
| SB | Adult Female | 416 | NA | 278 | 13 |
| SB | Juvenile | 169 | NA | 211 | 26 |

**Table S2:** The sample sizes used to generate $G_d$ for the PC.

| Parameter | Sample size |
|---|---|
| $a_{PC}$ | 42 |
| $a_{SB}$ | 2237 |
| $n_{PC}$ | 337 |
| $n_{SB}$ | 5101 |
